# Nucleotide usage biases distort inferences of the species tree

**DOI:** 10.1101/2020.07.30.228965

**Authors:** Rui Borges, Bastien Boussau, Gergely J. Szöllősi, Carolin Kosiol

## Abstract

Despite the importance of natural selection in species’ evolutionary history, phylogenetic methods that take into account population-level processes typically ignore selection. The assumption of neutrality is often based on the idea that selection occurs at a minority of loci in the genome and is unlikely to compromise phylogenetic inferences significantly. However, genome-wide processes like GC-bias and some variation segregating at the coding regions are known to evolve in the nearly neutral range. As we are now using genome-wide data to estimate species trees, it is natural to ask whether weak but pervasive selection is likely to blur species tree inferences. We developed a polymorphism-aware phylogenetic model tailored for measuring signatures of nucleotide usage biases to test the impact of selection in the species tree. Our analyses indicate that while the inferred relationships among species are not significantly compromised, the genetic distances are systematically underestimated in a node-height dependent manner: i.e., the deeper nodes tend to be more underestimated than the shallow ones. Such biases have implications for molecular dating. We dated the evolutionary history of 30 worldwide fruit fly populations, and we found signatures of GC-bias considerably affecting the estimated divergence times (up to 23%) in the neutral model. Our findings call for the need to account for selection when quantifying divergence or dating species evolution.

**Significance statement:** Although little is known about the impact of natural selection on species tree estimation, expectations are that it occurs at a minority of loci in eukaryotic genomes and is thus unlikely to affect the divergence process. However, growing evidence suggests that a large amount of the genomic variation evolves under weak but pervasive selection (e.g., fixation biases created by GC-bias gene conversion). We tested the impact of unaccounted-for nearly neutral selection on species tree estimation and found that the estimated branch lengths are systematically biased. Our results highlight the need for selection-aware models in species tree estimation and molecular dating.

## Introduction

Species trees provide a framework to understand the divergence process and speciation and are nowadays used routinely in integrative research to address many biological questions. Despite their generalized use, modeling species evolution using DNA sequences poses significant challenges. The main one being that different genes or genomic regions narrate different evolutionary histories, leading to discordant gene and species topologies (Maddison and Knowles, 2006). Incomplete lineage sorting (ILS), i.e., the maintenance of ancestral polymorphisms due to random genetic drift, is a primary cause of such discordance (Pollard et al., 2006). However, other processes were also described, such as gene gain or loss, horizontal gene transfer across the species boundaries, and gene flow among diverging populations (reviewed in Szöllősi et al. (2015)). Apart from the gene/species tree discordance, difficulties arise when quantifying divergence. Xu and Yang (2016) reported that calendar times and evolutionary rates can be particularly challenging to untie for deeper evolutionary scales where the molecular clock is violated.

Despite the challenges posed by modeling species evolution, the last two decades have seen an explosion of sophisticated statistical methods for inferring species trees. The multispecies coalescent model (MSC) has arisen as a leading framework for inferring species phylogenies while accounting for ILS and gene tree-species tree conflict (Rannala and Yang, 2003). The coalescent traces the genealogical history of a sample of sequences from a population backward in time, describing the stochastic process of lineage joining (Kingman, 1982a,b).

Nevertheless, alternative approaches to the MSC have been proposed, as the polymorphismaware phylogenetic models (PoMos; fig. 1) (De Maio et al., 2013, 2015). PoMos model the allele content of a set of populations over time at a particular locus, thus naturally integrating over all the possible allelic histories to directly estimate the species tree. (To be technically accurate, PoMos estimate population trees. However, because of PoMo ability to model closely related populations as well as diverged populations it most prominent application are populations sampled to estimate species trees. Therefore, we will adhere to this terminology in this article.) PoMos naturally account for ILS while avoiding using genealogy samplers that are computationally costly. Different from the MSC, PoMos assume independence between sites, which allows easily gathering information from multiple individuals and several populations to testing hypotheses genome-wide (Schrempf et al., 2016, 2019). More importantly, PoMos are versatile, for they can be straightforwardly rebuilt to account for other population forces (e.g., allelic selection (Borges et al., 2019)). MSC-based methods with selection are notoriously difficult.

**Figure 1:**
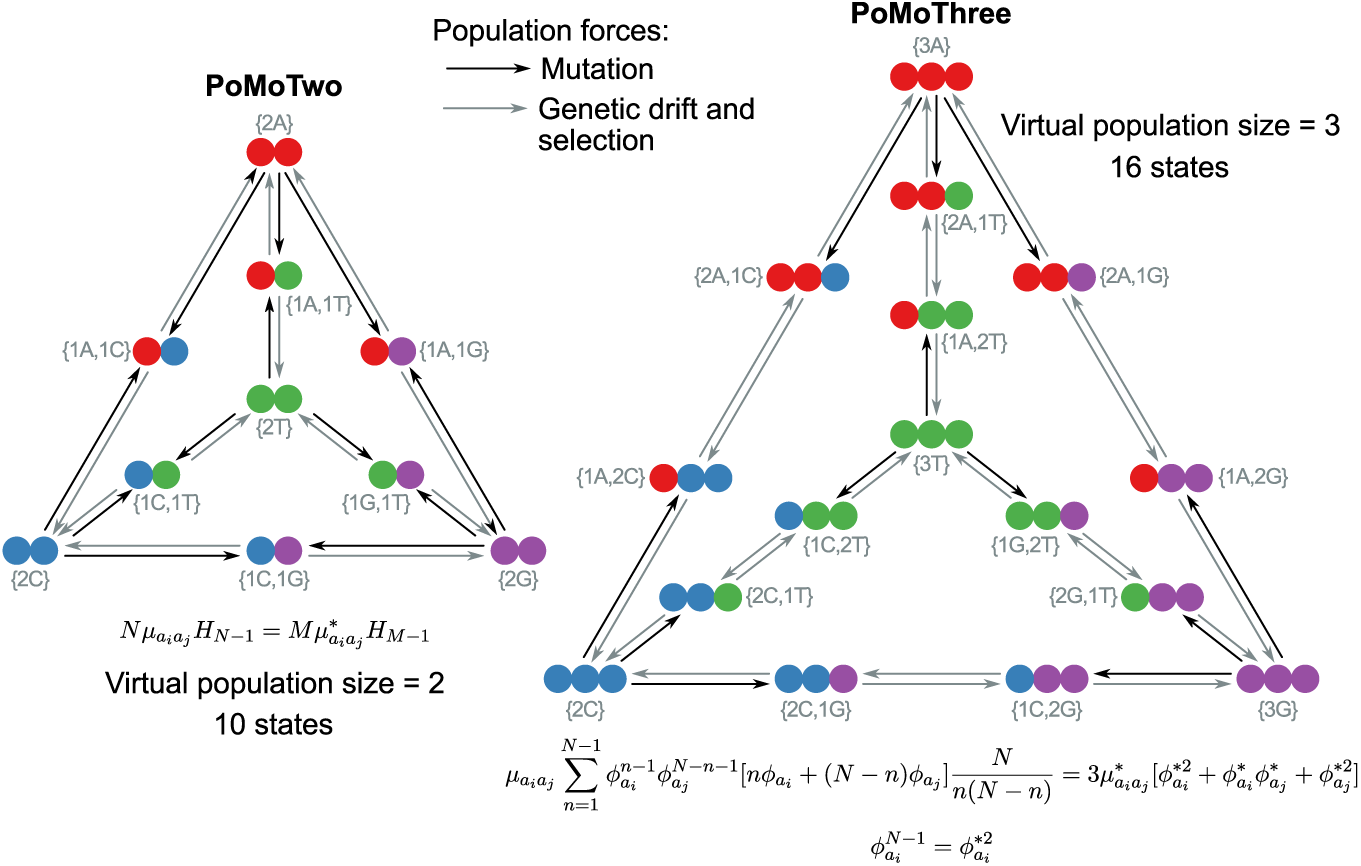
State-space and parameter scaling of the virtual PoMos. The tetrahedrons represent the state-space of the virtual PoMos: the fixed sites {*Ma*_*i*_} are placed in the vertices, while the edges represent the polymorphic states {*ma*_*i*_, (*M* − *m*)*a*_*j*_}. Black and grey arrows distinguish respectively mutation from genetic drift plus selection events. PoMo is a particular case of the four-variate Moran model with boundary mutations and selection, in which the alleles are the four nucleotide bases. The formulas represent the parameter scaling between the effective and the virtual populations (supplementary text S1). 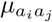 is the mutation rate from allele *a*_*i*_ to *a*_*j*_; and 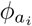 is the fitness coefficient of allele *a*_*i*_ on the effective dynamic of population size *N* (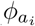 relates to the selection coefficient of *a*_*i*_ by the equation 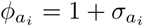). 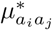 and 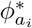 are the corresponding parameters in the virtual dynamic.

Despite the remarkable effort made to account for forces that can detail the species evolutionary history, models of species evolution generally ignore natural selection. While the literature acknowledges the need for selection-aware models of species evolution, the expectation is that most forms of natural selection are not greatly compromising phylogenetic analysis (Edwards, 2009). Two main arguments justify this expectation. The first argument is that selection often occurs at a minority of loci in the genome, in which case any spurious signals would likely be swamped out by the many neutral loci that are sampled (Edwards et al., 2016). The second argument is that the forms of selection that most likely affect the species tree are rare. This is the case of selection-driven convergent evolution and balancing selection, which tend to preserve beneficial alleles at a gene for long periods of time (Castoe et al., 2009; Edwards, 2009). As corroboration for these expectations, a recent study found that species-specific positive selection has mild effects on phylogeny estimation across an extensive range of conditions encountered in empirical data sets (Adams et al., 2018).

However, such claims have not been properly tested, especially for the case of pervasive weak selection. Theoretical expectations predict the existence of an intermediate category between the neutral and the selected modes of evolution, known as nearly neutral. The study of variants with small selection coefficients, particularly slightly deleterious mutations, has been the focus of the so-called nearly neutral theory (Ohta, 1973). This theory provided criteria to define the fate of nearly neutral variants, which depends on both selection and random genetic drift. Ohta (2002) has suggested that if the relative advantage or disadvantage, *σ*, of a particular allele is less than twice the reciprocal of the effective population size *N* (i.e., the scaled selection coefficient *N* |*σ*| < 2) the allele’s trajectory is nearly neutral. However, more lenient thresholds were also reasoned (e.g., Nei (2005)).

Empirical evidence for nearly neutral evolution has become more substantial within the last few years due to the possibility of sequencing many genomes and multiple individuals. One such example is GC-biased gene conversion (gBGC), a mutation bias favoring G and C alleles over A and T alleles at mismatch positions during recombination (Galtier et al., 2001). gBGC affects the fixation probability of GC alleles and is best modeled as selection (Nagylaki, 1983). Integrative analysis considering the recombination landscape and nucleotide substitution patterns along the genome provided evidence that gBGC acts in both eukaryotes and bacteria (Galtier et al., 2009; Lassalle et al., 2015). In humans, gBGC was estimated one order of magnitude lower than the reciprocal of the effective population size (∼ 1.17 × 10^−5^) (Glémin, 2010), thus, gBGC produces patterns of nucleotide substitution that resemble weak selection. The same was observed for other apes where most of their exons evolve under weak gBGC (*Nγ* < 1; where *γ* is the GC-bias rate) (Lartillot, 2013). In fruit flies, most amino acid replacements have weak signatures of positive selection. However, most of the selective effects are nearly neutral, with around 46% of amino acid replacements exhibiting scaled selection coefficients lower than two and 84% lower than four (Sawyer et al., 2007). Also, many non-synonymous polymorphisms in functionally important sites of human and bacterial populations were shown to segregate at frequencies of around 1–10%; i.e., much more frequent than variants associated with classic Mendelian diseases (Hughes et al., 2003; Hughes, 2005). Such an observation suggests that ongoing purifying selection among segregating amino acids is weak.

As a substantial part of the genome evolves nearly neutrally, this form of weak but pervasive selection has the potential to impact species’ evolutionary history. Here, we present novel PoMos specially tailored to measure nucleotide usage bias throughout the genome and use them to assess the impact of unaccounted-for weak selection (or selection-like processes) on species tree estimation. We show that near neutrality significantly blurs inferences of the species tree by biasing estimations of the species divergence and discuss the implications of these biases on molecular dating.

## Results

### Modelling species evolution with selection

PoMos offer a versatile approach to describe species evolution (Leaché and Oaks, 2017). They add a new layer of complexity to the standard models of sequence evolution by accounting for population-level processes (such as mutations, genetic drift, and selection) to describe sequence evolution. To do that, PoMos expand the 4 × 4 state-space to model a population of individuals in which changes in allele content and frequency are both possible. The PoMo state-space includes fixed (or boundary) states {*Na*_*i*_}, in which all *N* individuals have the same allele *a*_*i*_ ∈ {*A, C, G, T*}, but also polymorphic states {*na*_*i*_, (*N* − *n*)*a*_*j*_}, if two alleles *a*_*i*_ and *a*_*j*_ are present in the population with absolute frequencies *n* and *N* − *n* (fig. 1). PoMos do not consider triallelic sites, a simplification that is generally acceptable for eukaryotes, for which the levels of polymorphism are low (Lynch et al., 2016). Furthermore, like many phylogenetic methods, PoMos assume that sites are independent (i.e., linkage equilibrium).

One of the main challenges of PoMos is the size of their state-space grows with the effective population size: i.e., 4 + 6(*N* − 1) states. This prohibits inferences with realistic population sizes as they can easily have 10^4^ − 10^6^ individuals (Lynch et al., 2016), PoMos would typically employ 10 individuals because it results in manageable state-space and robust estimates of population properties such as the genetic diversity *θ* = 4*Nµ* (De Maio et al., 2015; Schrempf et al., 2016). Recently, we developed theory that permitted us to distinguish between the effective and virtual population size (Borges et al., 2019). Here, we use those results to develop the virtual PoMos. The idea is to mimic a population dynamic that unfolds on the effective population *N*, using a virtual population of smaller size *M*. It can be shown that by matching the expected diversity (i.e., the proportion of fixed and polymorphic sites) in both the effective and the virtual populations (Borges et al., 2019), one can obtain scaling laws for the mutation rates and fitness coefficients (supplementary Text S1 and fig. 1):

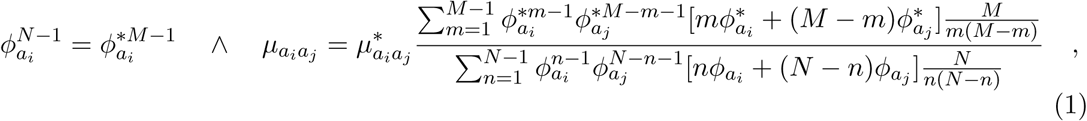

where 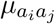 is the mutation rate from allele *a*_*i*_ to *a*_*j*_ and 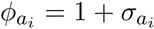 the fitness coefficient of allele *a*_*i*_ on the effective dynamic, while 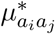 and 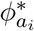 are the corresponding parameters in the virtual one.

Here, we defined the virtual PoMos Two and Three, by setting the virtual population size *M* to two and three individuals, respectively. PoMoTwo has only one polymorphic state per allelic pair, i.e., {1*a*_*i*_, 1*a*_*j*_}, and is used to describe species evolution under neutrality (fig. 1) while still accounting for mutation bias. For the neutral case, the mutation rates are scaled by the ratio of harmonic numbers (denoted here by *H*.) of the virtual and effective population sizes:

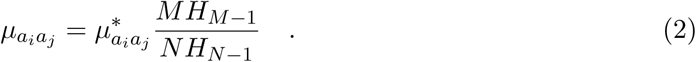

PoMoThree includes two polymorphic states per allelic pair, i..e, {1*a*_*i*_, 2*a*_*j*_} and {2*a*_*i*_, 1*a*_*j*_}, and additionally allows one to infer allelic selection (at least two polymorphic states are necessary to make the selection coefficients identifiable; fig. 1). The scalings in equation (1) simplify to:

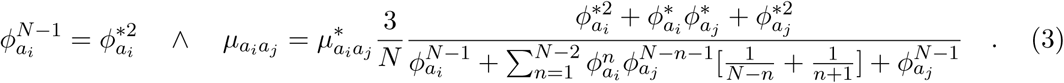

The virtual PoMoThree, in particular, allows quantifying signatures of selection and its impact on evolution.

### Virtual population dynamics provide a good fit for their effective counterpart

To validate the concept behind the virtual PoMos, we assessed their fit and computational performance using simulated data. We simulated phylogenetic data sets with signatures of weak but pervasive nucleotide usage biases as observed in populations of great apes and fruit flies (Borges et al., 2019; Prado-Martinez et al., 2013; Begun et al., 2007). In particular, we simulated the typically observed mutation bias from G or C (also known as strong alleles) to A and T (also known as weak alleles) opposing gBGC (Glémin et al., 2015).

Mutation rates were assumed reversible, where the mutation rate is defined by the product of an exchangeability term and an allele frequency under no selection: e.g., the mutation rate from a strong (S) to a weak (W) allele is defined as *µ*_*SW*_ = *ρ*_*SW*_ *π*_*W*_. The four possible mutation rates are thus fully determined by three parameters: *π*_*W*_ = 1 − *π*_*S*_, *ρ*_*SW*_ = *ρ*_*W S*_ and *ρ*_*SS*_ = *ρ*_*W W*_. The GC-bias preference was modeled by assigning a fitness of 1 to the weak alleles and a fitness *ϕ*_*S*_ = 1+*γ* to the strong alleles. A four-allelic Moran model was employed to simulate the evolutionary history of four populations across eight evolutionary scenarios by combining dynamics with high and low genetic diversity, weak and strong regimes of GC-bias, and shallow and deep divergence times. The specific parameters used in each simulated scenario are listed in table 1.

**Table 1.** Description of the simulated scenarios. *θ* represents the genetic diversity, *γ* the GC-bias coefficient (which relates to the GC-bias fitness by *ϕ*_*S*_ = 1 + *γ*) and *τ* the time of the most recent common ancestor in the phylogeny (defined in units of generations).

| Scenario | $\theta$ | $N\gamma$ | $\tau$ |
| --- | --- | --- | --- |
| 1 | 0.02 | 0.5 | $N$ |
| 2 | 0.02 | 0.5 | $10N$ |
| 3 | 0.02 | 1.5 | $N$ |
| 4 | 0.02 | 1.5 | $10N$ |
| 5 | 0.002 | 0.5 | $N$ |
| 6 | 0.002 | 0.5 | $10N$ |
| 7 | 0.002 | 1.5 | $N$ |
| 8 | 0.002 | 1.5 | $10N$ |

Phylogenetic data from the eight simulated scenarios and the virtual PoMoThree model were employed to infer the mutation rates and the GC-bias fitness on the virtual population. These estimates were recalibrated back to the effective dynamic using equation (1), and the relative bias was calculated to evaluate their accuracy. The GC-bias fitness shows the highest accuracy, being estimated with a consistent relative error smaller than 1% across the eight simulated scenarios. This result demonstrates that PoMoThree is particularly suited to quantify selection bias on genomic sequences. For most of the simulated scenarios, the mutation rate parameters are reasonably well estimated, with an average relative error smaller than 5% (fig. 2). These analyses indicate that the virtual PoMos provide a good approximation for an effective dynamic with mutation and selection biases. We have also estimated the mutation rate parameters using the virtual PoMoTwo and observed that the equilibrium base composition of strong alleles is systematically overestimated (i.e., *π*_*S*_ = 1 − *π*_*W*_), particularly in scenarios where GC-bias is stronger (supplementary fig. S1). This shows that the mutation rates try to accommodate the effect of selection towards the strong alleles by increasing the rate of weak to strong mutations (notice that *µ*_*W S*_ = *ρ*_*SW*_ *π*_*S*_).

**Figure 2:**
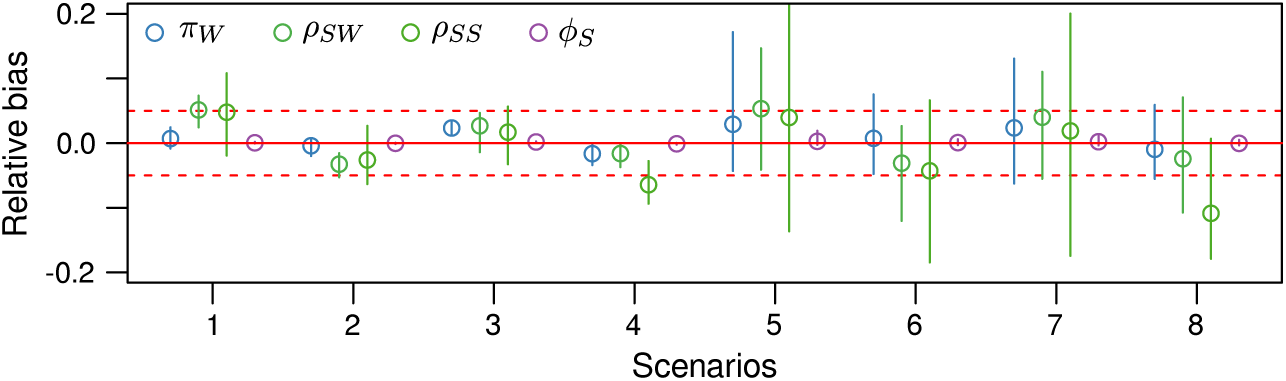
Estimating mutation rates and selection coefficients with a virtual dynamic. The simulated data from a four-allelic Moran dynamic with mutation bias and selection was fed to the virtual PoMoThree and the population parameters estimated using the Bayesian phylogenetic tool RevBayes (Höhna et al., 2016). As the population parameters were estimated in a population of three virtual individuals, we recalibrated them back to the effective population via the parameter scalings in equation (1). The estimated parameters are four: *π*_*W*_, *ρ*_*SW*_ and *ρ*_*SS*_ are terms of the mutation rates that were assumed reversible (i.e., *µ*_*SW*_ = *ρ*_*SW*_ *π*_*W*_); *ϕ*_*S*_ is the fitness coefficient of the strong alleles (i.e., the GC-bias fitness). The average bias of the estimates was obtained based on alignments of 10^5^ sites and 25 replicates per scenario. The specific parameters used in each simulated scenario are listed in table 1. The red dotted line represents a 5% relative error threshold.

We further compared the computational efficiency of the virtual PoMos, by comparing it to the standard general time-reversible nucleotide substitution model (GTR; Tavaré (1986)) in multiple sequence alignments, including 10 to 100 species and 100 to 1 million sites. For the three methods, similar settings for the phylogenetic inferences were employed in RevBayes (Höhna et al., 2016), including a standard MCMC chain with 50 000 generations and no parallelization. The computational performance of these three methods was assessed using three criteria: memory usage, CPU time, and average effective sample size of the sampled posterior parameters (ESS; i.e., the number of effectively independent draws from the posterior distribution that the Markov chain is equivalent to).

Despite PoMoThree and Two having a larger state-space (16 and 10 states respectively) and therefore more possibilities for different site patterns at the tips, the memory usage is relatively similar among the methods (fig. 3A). The memory usage of PoMoThree is on average 16% higher than that of PoMoTwo and 39% higher than that of GTR. However, we observed that in some cases, RevBayes could not start the analyses because the memory reached the computers’ maximum. Thus, while memory usage does not differ greatly among the methods, it certainly might be a limitation for genome-wide applications. Different from memory usage, the CPU time varies considerably among methods. We observed that the time consumed by each method increases exponentially with the alignment size (fig. 3B). PoMoThree requires, on average, 10.5 times more CPU time than the GTR and 2.3 times than PoMoTwo. This is because the PoMoThree has a larger state-space and consequently more site patterns to evaluate the likelihood. Regarding the average ESS, PoMoThree is the slowest mixing method with an average ESS of 2000 (supplementary table S1), 21.9% and 17.6% smaller than the average ESS of GTR and PoMoTwo, respectively.

**Figure 3:**
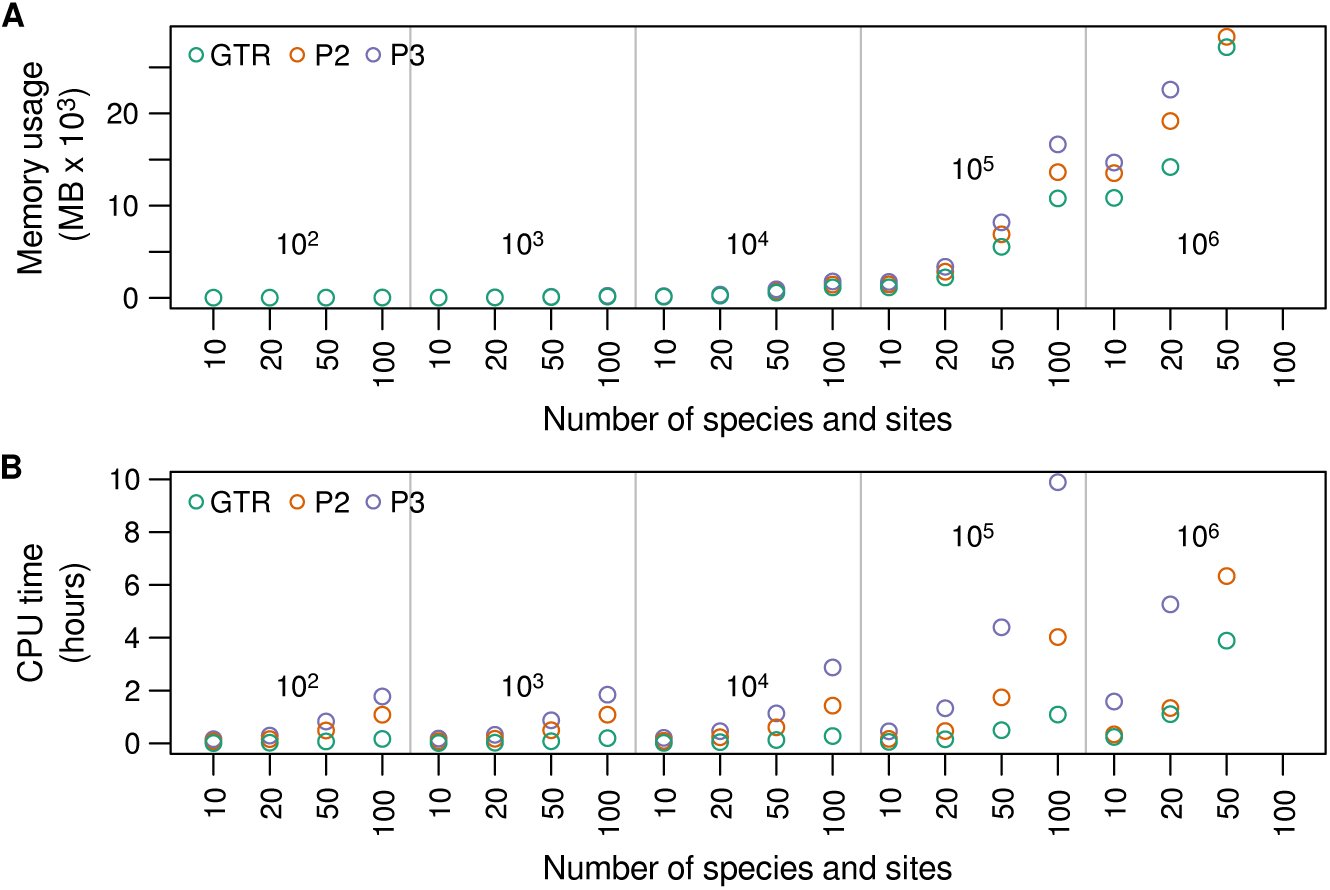
Computational performance of the virtual PoMos. The computational performance of the virtual PoMos Two and Three was compared with the general time-reversible model (GTR) based on **(A)** memory usage and **(B)** CPU time. The Bayesian inferences consisted of a standard MCMC chain with 50 000 generations and on simulated alignments with 10 to 100 species and 100 to 1 million sites, with levels of diversity similar to those of great apes (using the parameters defined for the simulated scenario 5). These analyses ran on an iMac desktop with an Intel Core i5 processor of 3.4 GHz of speed and 32GB of memory, without any kind of parallelization. Missing dots represent situations where the computer’s maximum memory was reached, and RevBayes was not able to start the analyses.

### Ignored selection affects species tree topology mildly

The impact of unaccounted-for selection on the reconstructed species tree can only be truly known by assessing the correctness of the estimation of both the topology and the branch lengths. Such aspects can be easily tested using simulated data on a known species tree. We used a species tree with four evolving populations, including two closely related populations and a third one recently diverging from the other two: `(A,(B,(C,D)))`. We set the evolutionary distances matching so that the first, second, and third divergence times are equal to *τ, τ*/2 and *τ*/4 respectively, where *τ* is the time of the most recent common ancestor. We thus expect higher levels of ILS when *τ* is equal to *N* generations than when it is equal to 10*N* (table 1).

To investigate the impact of selection on the species tree topology, we compared the trees inferred by the virtual PoMos (i.e., neutral PoMoTwo and the selection-aware PoMoThree) and the standard GTR model. If not stated otherwise, fixed and polymorphic counts from a sampled 10 individuals per population was used to perform the phylogenetic inferences with the virtual PoMos. As the GTR model only includes fixed states, a single random individual was used. The posterior probability of the correct topology was used to evaluate the accuracy of each method. Our results show that the virtual PoMos are more accurate than the GTR in all the tested scenarios, especially for alignments with fewer sites (fig. 4 and supplementary fig. S2). Because there are 15 different topologies, the expected posterior probability of the true tree if there is no signal in the data is around 6.7%, which is close to the performance of our methods in some conditions when there is little data. An interesting feature is that the topologies estimated by the GTR model tend to pick the first diverging branch incorrectly with greater probability than PoMos, especially for polymorphism-poor data sets (scenarios 1 and 5; fig. 4). This suggests that apart from being able to correct for ILS among closely related taxa, the PoMos are also superior in establishing deeper relationships. Despite that, we observed that the GTR’s accuracy increases with the alignment size, showing that the GTR topologies should converge to the correct species tree topology with genome-scale data.

**Figure 4.**
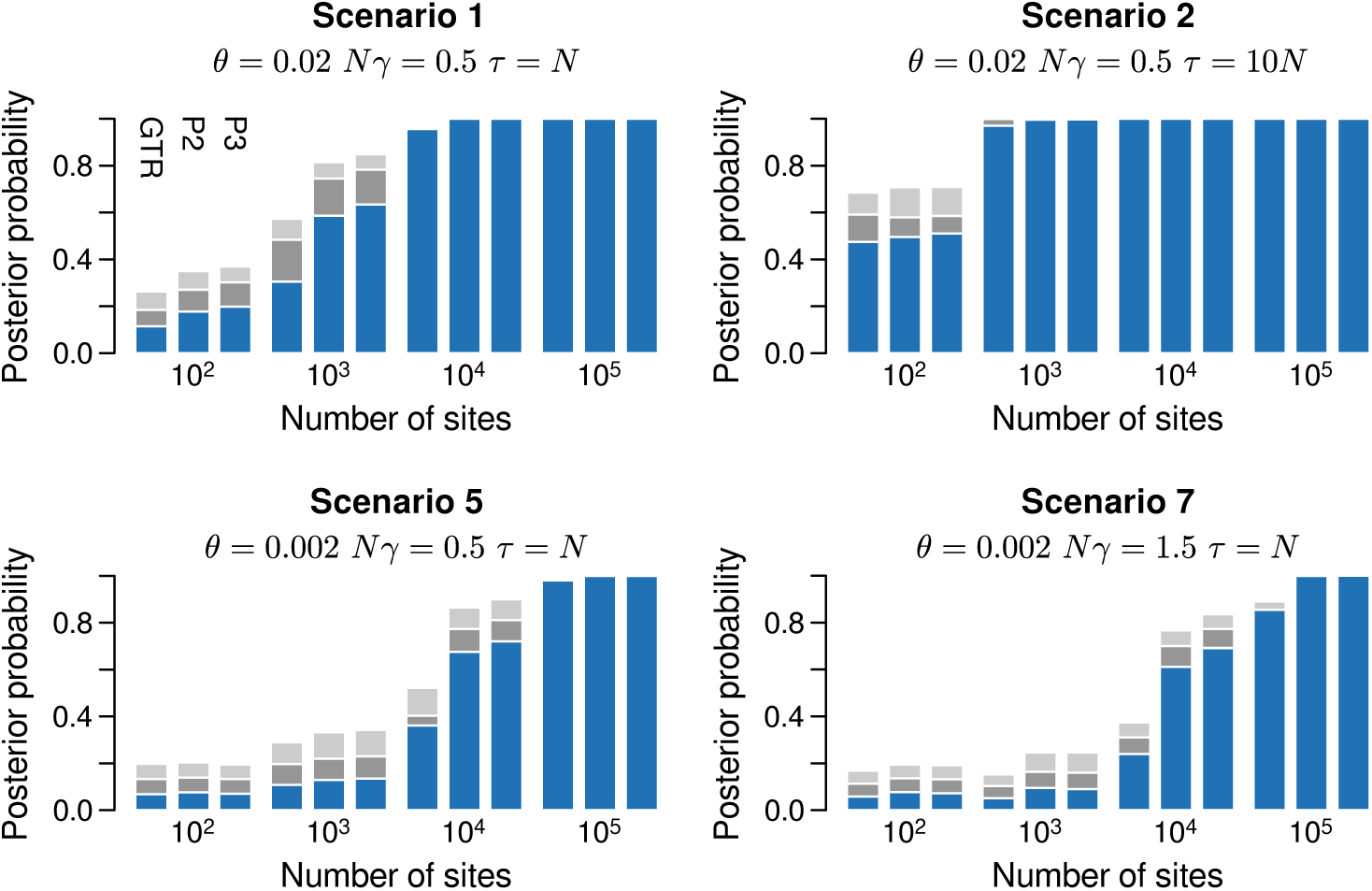
Estimating the species tree topology with selection. Data sets of different sizes (x-axis) and simulated under different scenarios were used to perform species tree inferences with three methods: GTR, PoMoTwo (P2 in the figure), and PoMoThree (P3 in the figure). We investigated the accuracy of the topology by assessing the posterior probability of the true topology (`(A,(B,(C,D)))`; blue sub-bars) under these three methods. The remaining probability corresponds to wrongly estimated topologies. The posterior probabilities of wrong topologies that are still able to correctly identify the first taxon to diverge are represented in shades of gray: `(A,(C,(B,D)))` in dark gray and `(A,(D,(C,B)))` in light gray. PoMoTwo and Three trees were inferred based on a sample of 10 individuals per taxa (totaling 40 sequences). The results for the eight scenarios can be found in supplementary fig. S2.

PoMoThree is only slightly more accurate than PoMoTwo, which indicates that neutral models may already provide fairly good estimates of the species tree topology (fig. 4). This is still valid when we increase the intensity of GC-selection (scenario 5 vs. 7; fig. 4)), suggesting that weak selection regimes should not greatly impact inferences of the relationships among species. Overall, our results confirm that the three methods perform similarly in estimating the species tree topology if enough genomic data is available. The specific number of sites necessary to return the correct topology will depend on several aspects, as the number of taxa and the complexity of the phylogenetic relationships among them. In any case, our results suggest that regardless of the employed method, better estimates of the species tree topology are expected for well-diverged taxa and polymorphism-rich data sets (scenario 1 vs. 2 and 7; fig. 4).

Similar conclusions can be drawn from when a sample of two individuals per population is taken instead of 10. Though the overall accuracy of PoMos decreases (supplementary fig. S3), we again observed that the posterior probability of the correct topology increases with the alignment size. This result is not surprising if we consider the results we have already discussed for the GTR model. Choosing a single virtual sequence to represent the whole population, as is often done when employing the standard nucleotide models with multi-individual data, corresponds to sample a single individual for the population (i.e., the situation where no polymorphisms are observed). Nevertheless, despite the dramatic reduction of the observed diversity, we observed that the GTR model can estimate the correct topology with enough genomic data.

### Neglected selection distorts the measured divergence

We investigated the impact of selection on the branch lengths by comparing the genetic distances estimated by the virtual PoMos and the standard GTR model. To assess the accuracy of the branch length estimates, we evaluated the absolute error between the estimated and expected pairwise distances among the four taxa. This strategy has the advantage of permitting comparing methods even if the estimated topologies are not exactly matching. Normalized distances were used for this comparison, as the GTR and PoMos operate in different evolutionary units: number of substitutions and Moran events (i.e., mutations plus frequency shifts), respectively.

Our results show that the error in the estimated branch lengths decreases as the size of the alignments increases (fig. 5 and supplementary fig. S4). Again, we observed that the GTR is the model of evolution producing the least accurate estimates, with an average error 2.5 times higher than PoMoThree, indicating that polymorphisms have a considerable impact on branch length estimation. PoMoThree outperforms PoMoTwo for all the observed scenarios, with PoMoTwo having an increased error of 61% on average. In some scenarios, PoMoThree has a negligible error (e.g., scenarios 2 and 8; Fig. 5 and supplementary figs. S4 and S5), demonstrating that the virtual PoMos provide a good fit for the divergence process operating in an effective dynamic with mutation and selection biases. If we only consider the scenarios where selection is more intense (i.e., *Nγ* = 1.5), the error increases to 75%, indicating that ignored selection significantly contributes to bias the estimated divergence (e.g., scenarios 6 vs. 8; Fig. 5 and supplementary figs. S4 and S5).

**Figure 5:**
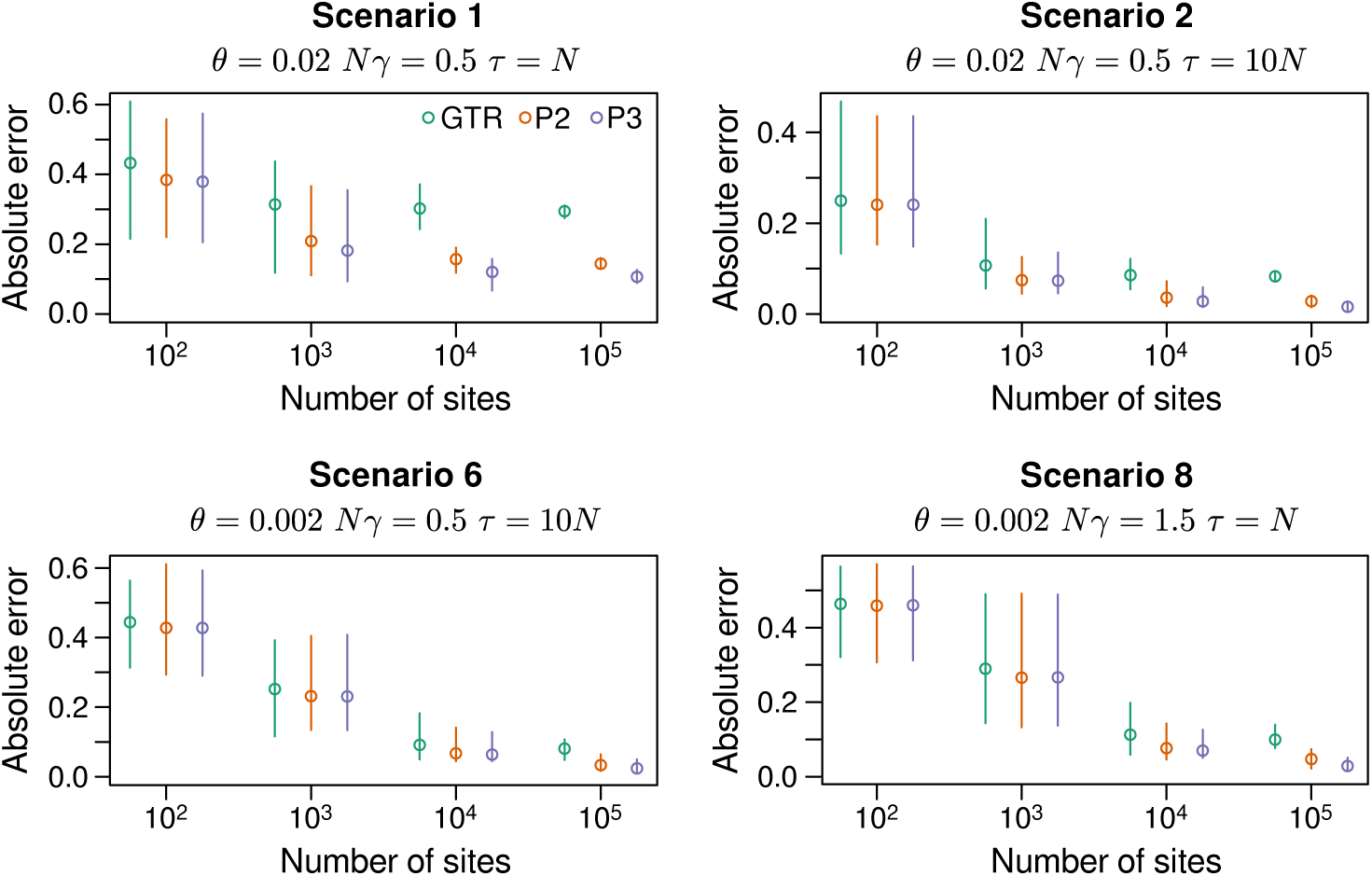
Estimating the species tree branch lengths with selection. Data sets of different sizes (x-axis) and simulated under different scenarios were used to perform species tree inferences with three methods: GTR, PoMoTwo (P2 in the figure), and PoMoThree (P3 in the figure). We investigated the accuracy of the branch lengths by calculating the absolute error of all six pairwise comparisons of genetic distances among the four taxa. As the three methods employ different evolutionary units, the branch lengths were normalized so that the three models could be compared. PoMoTwo and Three trees were inferred based on a sample of 10 individuals per taxa (totaling 40 sequences).

We have also observed that the error on the estimated branch lengths does not decay to zero as fast as for the topology case, persisting in some cases even when the topologies are perfectly estimated (inferences with 10^5^ sites in scenarios 1 and 7; fig. 5). Because of the sampling of individuals, polymorphisms in the original population may appear as fixed differences and provide misleading phylogenetic signal that could contribute to this slow decay. We observed that decreasing the sample size from 10 to two individuals increases the error of the estimated branch lengths by 75% on average in the virtual PoMos (supplementary fig. S5). As the patterns of diversity on the original and sampled populations are different, the models of evolution are unable to determine the exact contribution of mutations, fixations, and frequency shifts for the divergence process. The impact of such sampling bias should be particularly problematic among recently diverged populations, where several populations likely share segregating polymorphisms. Indeed, our simulations corroborate this expectation: scenarios 1 vs. 2 (fig. 5).

To establish that the superior performance of PoMoThree was not due to a larger state space with more polymorphic states, we have calculated the accuracy of the estimated topologies and branch lengths by employing the PoMoThree model with selection coefficients fixed to 0.0. We observed that this neutral PoMoThree has an intermediate accuracy between PoMoTwo and PoMoThree with selection (supplementary figs. S6 and S7), which suggest that modeling selection does increase the fit of PoMos beyond that what would have been expected by having a finer state space with more polymorphic states.

To further investigate the source of bias on the branch lengths, we tested whether the placement of the estimated divergence varies across evolutionary scales. To do that, we compare the height of each node on the true tree as estimated by the neutral PoMoTwo and the selection-aware PoMoThree models. For this comparison, we have only used node heights estimated under the 100 000 sites alignments, where both methods estimate the true topology correctly. We observed that PoMoTwo produces smaller node heights on average than PoMoThree (clouds always below the identity line in fig. 6A). This result suggests that weak selection signatures can mislead neutral models to estimate species histories with more recent divergence events. However, and more importantly, this bias is not uniform across nodes. The shallow nodes are less underestimated than the deeper nodes, with the ratio of node heights moving away from 1.0 from the shallow to the deeper nodes in all the tested scenarios (fig. 6A and supplementary figs. S8 and S9). The rate of evolution is known to be challenging to untie at deeper evolutionary scales (Xu and Yang, 2016); here, we show that ignoring pervasive weak selection may contribute to it.

**Figure 6:**
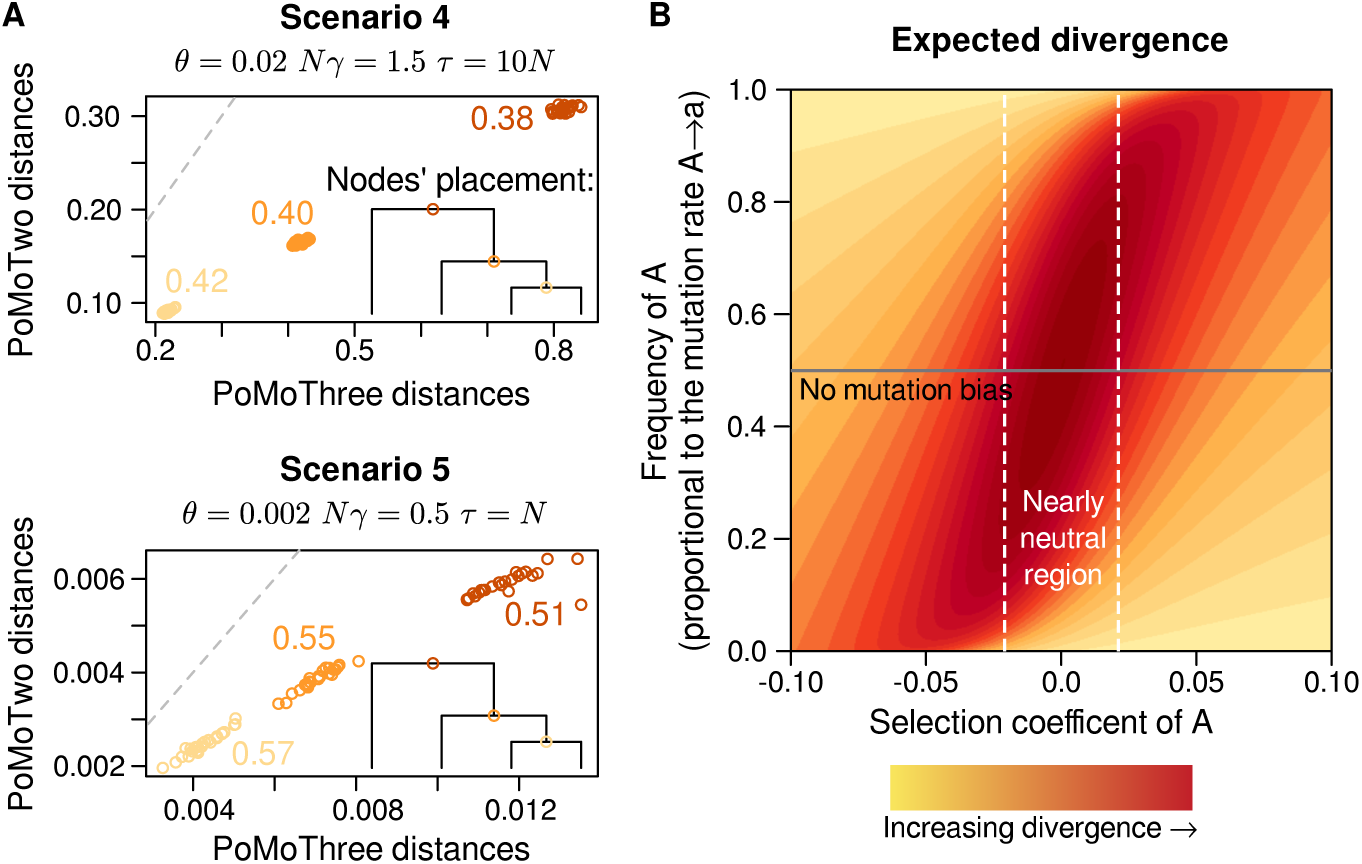
Estimating divergence in the presence of selection. **(A)** We investigated the association between the node heights as estimated by the virtual PoMos Two and Three. The phylogenetic tree depicts the three nodes’ placement in the true phylogeny: the node heights double as we approach the root. Values accompanying each cloud denote the average ratio of the node heights in PoMoTwo to those in PoMoThree; these ratios are always smaller than 1.0 for all the other simulated scenarios (supplementary figs. S8 and S9). Each cloud represents 25 replicated alignments. The depicted node heights were all estimated using the 100 000 alignments, where both methods return the correct topology with maximum accuracy. **(B)** The expected divergence was calculated for different regimes of selection and mutation. For the biallelic case, the base composition *π* is proportional to the mutation bias: 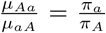, where *µ* is the mutation rate. If the base composition is equal (gray line), there is no mutation bias (supplementary text S2).

While our simulations intend to represent realistic scenarios, one might want to know what is the overall impact of selection (not necessarily nearly neutral) on the branch lengths. An advantage of PoMos is that some quantities can be formally obtained, such as the expected divergence per generation in diverse mutation-selection regimes (fig. 6B and supplementary text S2).

We observed that the expected divergence is generally higher for the neutral case and tends to decrease as selection becomes more intense (fig. 6B; such trend is not qualitatively affected by the effective population size as shown in supplementary fig. S10). Directional selection fades alleles that could potentially be drifting in the population to fixation, overall reducing the expected divergence: i.e., for the same amount of time, we expect more homogeneous sequences if directional selection is acting. Consequently, to match the same patterns of diversity, selection-aware models tend to return larger branch lengths than the neutral models, corroborating the results obtained in fig. 6A. Furthermore, the underlying population dynamic strongly determines the magnitude of biases on the estimated branch lengths. For instance, the expected divergence is generally higher when selection counteracts mutation (i.e., rarer alleles are favored and *vice versa*, as is the case with gBGC) than when selection and the mutational bias act concordantly (i.e., rarer alleles are disfavoured and *vice versa*; fig. 6B).

To assess the impact of weak selection on the species trees, we have here simulated and estimated genome-wide effects of GC preference. We note that this strategy is a simplification of the means by which gBGC acts, as it behaves transiently along the genome, being stronger close to recombination hotspots that rapidly change their location at phylogenetic scales (Duret and Galtier, 2009; Romiguier and Roux, 2017). Nevertheless, we note that our RevBayes implementation allows for setting more realistic scenarios, including branch-specific models or regional variation on GC-bias rates along the sequence alignment.

### Molecular dating with fruit fly populations

The geographic origins of the globally distributed fruit fly (*Drosophila melanogaster*) are still not fully understood. This is certainly not due to the lack of genomic data, as hundreds of sequenced genomes are available for the fruit flies (e.g., Hervas et al. (2017)), but primarily due to the methods of species tree estimation not scaling with multiple populations and genome-wide data. We employed the virtual PoMos to estimate and date the evolutionary history of 30 fruit fly populations. We used 966 sequences (supplementary table S2) from 1 million distantly located genomic sites, acknowledging the sites’ independence assumption. We also assumed a global clock for molecular dating analyses, which is generally valid for short time scales. We set a uniform prior of 60 000 ± 15 000 years dating the African population expansion, as is suggested in the field’s literature (e.g., Li and Stephan (2006); Stephan and Li (2007); Laurent et al. (2011)). The phylogenies estimated by the virtual PoMos agreed for most of the major clades (fig. 7), overall supporting a similar phylogeographic history for the fruit fly populations. Our analyses showed that the fruit fly populations have flourished in the southeast of the African continent. It is generally accepted that fruit flies originated in equatorial Africa (Lachaise et al., 1988; Stephan and Li, 2007). Our analyses suggested further that the species spread to the north, establishing in western equatorial Africa. From there, fruit flies colonized the western equatorial and the northern regions of Africa, with the northern wave leading to the spread of fruit flies worldwide. This phylogeography is not captured by the GTR model (supplementary fig. S11), probably due to high levels of ILS among fruit flies. Indeed, we found that 4.8% of polymorphisms are shared by three or more populations, and 2.0% by 10 or more populations.

**Figure 7:**
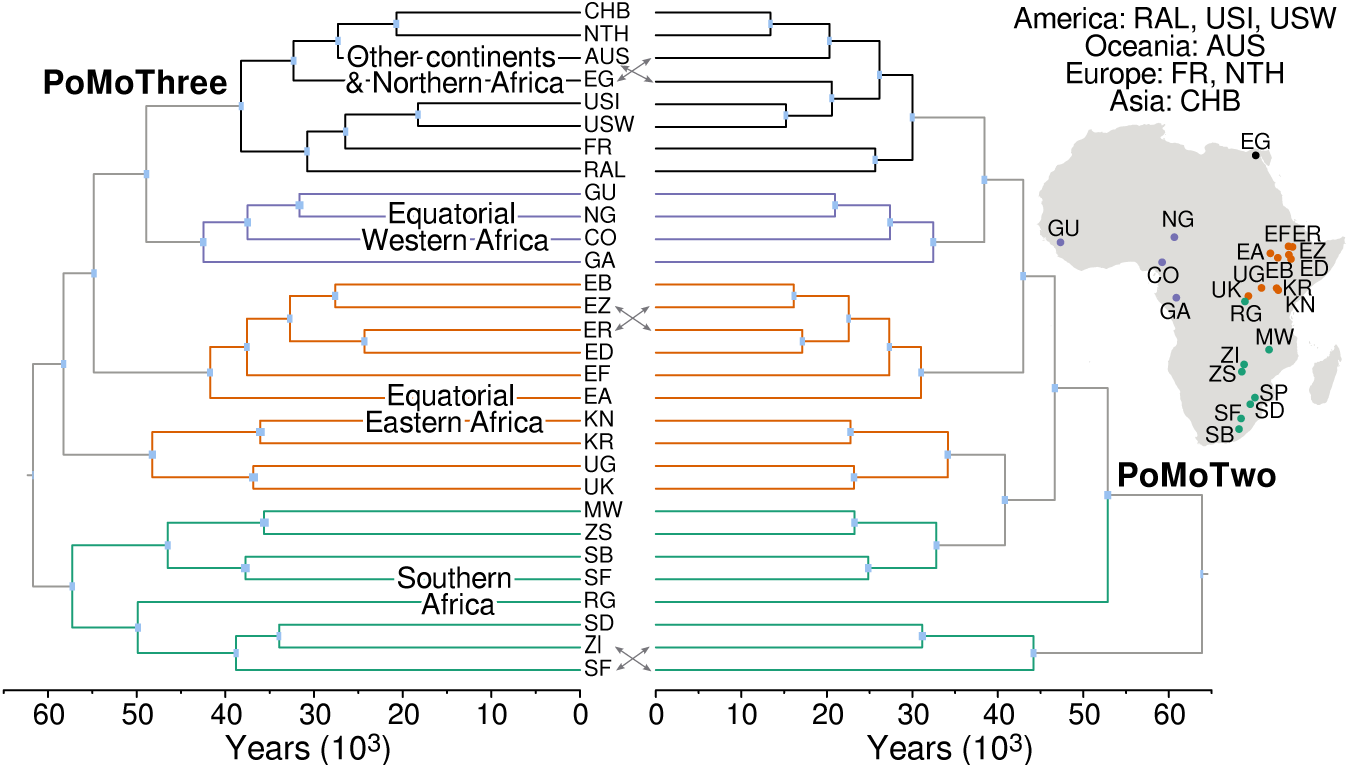
Dating the evolutionary history of 30 worldwide fruit fly populations. We dated the evolutionary history of 30 populations with 4 to 205 individuals (966 sequences in total, supplementary table S1) of *Drosophila melanogaster* distributed across the globe (data taken from Hervas et al. (2017)). The phylogenetic analyses were employed in RevBayes (Höhna et al., 2016), using alignments of 1 million distantly located genomic sites and the PoMoTwo and PoMoThree models of evolution, the latter being selection-aware. The gray lines between the trees indicate disagreement between the two models. The maximum *a posteriori* tree is depicted in both cases. The blue rectangles represent the 95% credibility intervals of the age of each node.

The time inferences are less consistent between the virtual PoMos, with PoMoThree providing, as expected by our formal results and simulations, more ancient node ages (around 23%; figs. 7 and supplementary figs. S12 and S13). Our divergence times are also comparatively more ancient than other studies. For example, some studies suggested a split of the African lineage at about 16 000 years ago (Stephan and Li, 2007; Laurent et al., 2011), while our analyses estimated it around 38 000 years ago. While unaccounted signatures of selection can underestimate the divergence times, we also notice that the non-monophyly of the out-of-Africa fruit fly populations (i.e., the Egyptian population (EG in fig. 7) clades together with these populations). This can cause such volatility in the estimated divergence times among studies. We note that the narrow time intervals obtained in our dating analyses should be taken with care, as we are using a sample size of one million sites and a while more complex still simplifying models of evolution and molecular clock. The influence of large data on the time divergence intervals is most clear with the GTR model, which infers a completely different phylogeny than PoMos but with quite accurate time intervals (supplementary fig. S11).

An additional advantage of using selection-aware methods is that we can quantify selection coefficients. We measured the fruit flies’ GC-bias rate at around 6.17 × 10^−7^ per site per generation, one order of magnitude lower than the reciprocal of their effective population size (approximately 1 million individuals according to Lynch et al. (2016)) and matching the expectations of gBGC. It remains unclear whether fruit flies have gBGC (Robinson et al., 2014). While it is difficult to think of any other process than gBGC favoring G and C alleles genome-wide, further studies associating these signatures with the recombination landscape are still needed.

## Discussion

Methods of species tree estimation have become an essential tool in studies of molecular evolution. Despite their widespread use, most of these methods assume neutrality. Here, we showed that unaccounted-for pervasive selection and selection-like processes such as gBGC significantly impact the inferences of the species tree. Our extensive analyses in both simulated and real data have two main implications for phylogenomic studies.

The first one is that assumed neutrality is likely to bias genetic distances in typical population genetic data sets. Modeling natural selection within the species evolution framework is not theoretically or computationally trivial, which has undoubtedly contributed to most tree inference methods making the neutral assumption. The virtual PoMos are full-likelihood methods that include selection and scale with genome-wide data from multiple populations and individuals. Other species tree inference methods, like the MSC, experience a data bottleneck for a few sequences and a handful of populations. Constraints between the species tree and the genealogical histories make it difficult to perform their joint inference. Furthermore, the space of unknown genealogies is large, and the genealogy samplers computationally intensive (Flouri et al., 2020; Rannala et al., 2020). In contrast, PoMos directly estimate the species tree and naturally integrate over all possible genealogies. Hence, we expect that our methods will permit us to understand better the practical consequences of assuming neutrality in real contexts. Here, we have focused on the case of weak pervasive selection for populations with patterns of diversity similar to fruit flies and great apes. However, our results are likely to extend to several other biological groups, as GC-biased gene conversion is found all across the major groups of eukaryotes (Pessia et al., 2012).

The second one is that the bias created by ignoring selection is not uniform across nodes; it is more significant in the deeper nodes than in the shallow ones. Previous reports reasoned that directional selection should only mildly affect the rate of evolution (Edwards, 2009). By demonstrating that this effect is node-height dependent, we show that merely recalibrating the overall expected divergence is insufficient to correct divergence time inferences. Therefore, selection-aware models are particularly crucial for molecular dating or other phylogenetic analyses relying on the estimated branch lengths. By showing that one cannot ignore the net effect of selection in the evolutionary process (even in its weakest forms as tested here), we question the neutral assumption’s suitability in species tree inference.

As the number of genomes sequenced increases rapidly, the development of methods describing the evolutionary process in all its complexity emerges as of fundamental importance. The methods presented here represent an important extension of existing approaches by allowing selection and permitting inference with large-scale data. In the future, we envision that our methods will be particularly useful for molecular dating using genome-wide data sets.

## Materials and Methods

### Simulations

As our methods aim at estimating species trees with genomic sequences, our simulations were designed to imitate general patterns of molecular evolution. In particular, we set a molecular dynamic with GC-bias favoring the fixation of G and C alleles (or strong alleles) and mutational bias to A and T alleles (or weak alleles) (Prado-Martinez et al., 2013; Begun et al., 2007). Mutation rates were assumed reversible and defined by the product of an exchangeability term and an allele frequency: e.g., the mutation rate from a strong to a weak allele is *µ*_*SW*_ = *ρ*_*SW*_ *π*_*W*_. Exchangeabilities between weak and strong alleles (i.e., *ρ*_*SW*_ = *ρ*_*W S*_) were set three times more likely than between similar alleles (i.e., *ρ*_*SS*_ = *ρ*_*W W*_). We have also set the frequency of the weak alleles to be higher than the strong alleles (i.e., *π*_*W*_ = 0.7 and *π*_*S*_ = 0.3). By combining the two exchangeabilities and allele frequencies, four mutation rates can be computed: *µ*_*SW*_ = *ρ*_*SW*_ *π*_*W*_, *µ*_*W S*_ = *ρ*_*SW*_ *π*_*S*_, *µ*_*SS*_ = *ρ*_*SS*_*π*_*S*_ and *µ*_*W W*_ = *ρ*_*SS*_*π*_*W*_. A fitness of *ϕ*_*S*_ = 1 + *γ* was set to the strong bases, where *γ* is the GC-bias rate, while the weak bases were assumed neutral with fitness 1.

We simulated the evolution of a population of four populations of 100 individuals using the PoMo transition matrix, which embedded Markov chain is equivalent to the four-variate Moran model with boundary mutations and allelic selection. The simulator was implemented in R (R Core Team, 2021) and is provide in the additional files (see data availability statement). The mutation rates and the selection coefficient were set according to the previously defined molecular dynamic and appropriately rescaled. To make our simulation more realistic, we defined different evolutionary scenarios. Following the patterns of diversity observed in populations of fruit flies and great apes (Prado-Martinez et al., 2013; Begun et al., 2007)), we varied the levels of genetic diversity by setting *θ* = 0.02 or 0.002. We added two regimes of GC-bias with scaled selection coefficient *Nγ* = 0.5 or 1.5, both under the nearly neutral range (i.e., *Nγ*< 2). We have also considered different divergence times by setting the time of the most recent common ancestor to *τ* = *N* or 10*N* generations. The evolution of four populations was simulated according to the following phylogenetic tree `(((pop4:`*τ*/4`, pop3:`*τ*/4):*τ*/4`, pop2:`*τ*/2):*τ*/2`, pop1:`*τ* );. As some populations are closely related, we expect considerable levels of ILS for shorter divergence times. The simulated scenarios and respective parameters are listed in table 1. For each scenario, we produced alignments of 100 to 10^5^ sites and 25 repetitions. We further generated two additional data sets by sampling 10 and 2 individuals per population. To assess the computational performance of the virtual PoMos, we simulated phylogenetic data for 10 to 100 populations and alignments of 100 to 1 million sites based on the parameters of scenario 5.

Because the simulations were performed in terms of the effective dynamic, the simulated data was converted to match the GTR, PoMoTwo, and PoMoThree state spaces. The fixed states in the original population {*Na*_*i*_} were directly converted to the fixed states {3*a*_*i*_} and {2*a*_*i*_} in the virtual PoMoThree and Two respectively. The polymorphic state {*na*_*i*_, (*N* − *n*)*a*_*j*_} in the original population was directly converted to the state {1*a*_*i*_, 1*a*_*j*_} PoMoTwo state, as this model has only one polymorphic state per polymorphic type. Differently, PoMoThree has two polymorphic states per polymorphic type. In this case, we used the allele frequency 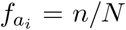 in the original state and attributed the state {2*a*_*i*_, 1*a*_*j*_} with probability 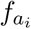 and {1*a*_*i*_,2*a*_*j*_} with probability 1 − 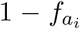. As the GTR state-space only includes the four alleles, fixed states were set to the fixed allele {*a*_*i*_} while polymorphic ones to one of the alleles, {*a*_*i*_} or {*a*_*j*_}, by randomly drawing according to the allele frequency. The state conversion for the GTR model corresponds to sample a single individual from the population.

### Bayesian phylogenetic inferences

Phylogenetic trees were estimated using Bayesian inference in the RevBayes software (Höhna et al., 2016). The virtual PoMos are implemented in RevBayes and can be employed using the functions `fnReversiblePoMoTwo4N` and `fnReversiblePoMoThree4N`, which can be found in the RevBayes GitHub project (https://github.com/revbayes). Alternatively, the general function `fnReversiblePoMo4N` can also be used by setting the population size to two or three. In total, four free parameters were estimated during the Bayesian inferences: *π*_*W*_, *ρ*_*W S*_, *ρ*_*SS*_ = *ρ*_*W W*_ and *ϕ*_*S*_. We set a Beta prior on *π*_*W*_ and an exponential prior on the exchangeabilities and the GC-bias fitness *ϕ*_*S*_. We assumed a strict global clock rate for the molecular clock, drawn from an exponential prior with a root age fixed to 1.0. We fixed the root age to allow distance comparisons between the GTR and the virtual PoMos, as they all operate with different evolutionary units. A uniform time tree prior was set for the phylogeny. MCMC estimation was carried out with two chains with 50 000 generations for the GTR model and 75 000 for the virtual PoMos. The ESS was checked for convergence. The MCMC samples were accepted provided that the ESS of all model parameters was above 200. Similar priors were used for the GTR model, with the exception that the exchangeabilites were also set with a beta prior instead of an exponential one.

### Application to fruit fly populations

Genome-wide data from 30 worldwide distributed populations of *Drosophila melanogaster* with 4 to 205 individuals (966 individuals) were extracted from the PopFly database (Hervas et al., 2017). We have only used genomic data from the 2R, 2L, 3R, and 3L autosomes. In total, one million genomic sites were randomly sampled for the phylogenetic inferences. The fruit flies allele counts were converted to the GTR, PoMoTwo, and Three state spaces following the same procedure as for the simulated data. The resulting multiple sequence alignments were then used to perform molecular dating analyses in RevBayes (Höhna et al., 2016). We employed a global clock model together with a uniform prior of 60 000 ± 15 000 years dating the African population expansion, following the results of Laurent et al. (2011). For the remaining parameters, we set the same priors as for the Bayesian phylogenetic inferences with the simulated data.

## Data Availability Statement

The simulated data, RevBayes scripts, and fruit fly alignments generated and analyzed during the current study are available at Zenodo (https://doi.org/10.5281/zenodo.5202827).

## Author Contributions

R.B., C.K., and G.S. designed research; R.B. and B.B. implemented the PoMo models in RevBayes; R.B. and C.K. analyzed data; R.B., C.K., G.S. and B.B. wrote and revised the paper.

## Acknowledgements

The molecular dating analyses of the fruit fly data set have been achieved using the Vienna Scientific Cluster (VSC). This work was supported by the Vienna Science and Technology Fund (WWTF) [MA16-061] and partially supported by the Austrian Science Fund (FWF) [P34524-B]. GJS received funding from the European Research Council under the European Union’s Horizon 2020 research and innovation program under grant agreement no. 714774 and the grant GINOP-2.3.2.-15-2016-00057. We thank two anonymous reviewers and the associate editor Mario dos Reis for their comments and suggestions, which helped us to improve the manuscript.

