## Supplementary material for "Nucleotide usage biases distort inferences of the species tree"

This PDF file includes:

- Text S1 and S2
- Figs. S1 to S6
- Tables S1 and S2
- References

#### Supplementary Text S1. Rate matrix and stationary distribution of PoMos

PoMos model the evolution of a population of $N$ individuals and $K$ alleles in which allele content and frequency changes occur. These are mediated by population forces, such as mutation, genetic drift, and selection. The PoMo state-space includes fixed (or boundary) states $\{Na_i\}$, in which all $N$ individuals have the same allele $i \in \{0, 1, ..., K-1\}$, and polymorphic states $\{na_i, (N-n)a_j\}$, in which two alleles $a_i$ and $a_j$ are present in the population with absolute frequencies $n$ and $N-n$.

Mutations occur with rate $\mu_{a_i a_j}$. Mutations govern the allele content and only occur in the fixed states:

$$q_{\{Na_i\} \to \{(N-1)a_i, 1a_j\}} = \mu_{a_i a_j} \quad . \tag{1}$$

Often, a reversible mutational model is considered. In this case, we break the mutations into a base composition $\pi$ and exchangeability parameter $\rho$ (i.e., $\mu_{a_i a_j} = \rho_{a_i a_j} \pi_{a_j}$) just like the GTR. However, in PoMos, we model mutations, not substitutions. Such an assumption can still model mutational biases quite well and simplifies obtaining formal quantities with PoMos.

Genetic drift is modeled according to the Moran model, in which one individual is chosen to die, and one individual is chosen to reproduce at each time step. Selection acts to (dis)favor alleles by differentiated fitnesses: $\phi_{a_i}$. Together, genetic drift and selection govern the allele frequency changes:

$$q_{\{na_i, (N-n)a_j\} \to \{(n+1)a_i, (N-n-1)a_j\}} = \frac{n(N-n)}{N[n\phi_{a_i} + (N-n)\phi_{a_j}]} \phi_{a_i} \quad . \tag{2}$$

We accelerate time by a factor of $N$ (i.e., multiply the rates by $N$), so that time is expressed in generations. Like the standard substitution models, PoMos are continuous-time Markov models and are fully characterized by their rate matrices. The rates in (1) and (2) define the PoMos rate matrices. If the mutation rates are reversible, the PoMo rate matrices enjoy reversibilitity. This property allow us to easily determine its stationary distribtution $\psi$ via the detailed balance equations. The stationary frequencies of the fixed states are defined by

$$\psi_{\{Na_i\}} = \pi_{a_i} \phi_{a_i}^{N-1} \quad , \tag{3}$$

while for the polymorphic states are

$$\psi_{\{na_i, (N-n)a_j\}} = \pi_{a_i} \pi_{a_j} \rho_{a_i a_j} \phi_{a_i}^{n-1} \psi_{a_j}^{N-n-1} [n\phi_{a_i} + (N-n)\phi_{a_j}] \frac{N}{n(N-n)} \quad . \tag{4}$$

The stationary distribution is very informative as it informs on the frequency of fixed and polymorphic states of the different allele types, overall characterizing populations' diversity. We define the virtual PoMos by considering the population dynamic operating in two populations of different sizes: $N$ and $M$ individuals. We then established their stationary distribution and equaled the frequencies of the fixed and polymorphic states: i.e.,

$$\psi_{\{Na_i\}} = \psi_{\{Ma_i\}} \quad \wedge \quad \sum_{n}^{N-1} \psi_{\{na_i, (N-n)a_j\}} = \sum_{m}^{M-1} \psi_{\{ma_i, (M-m)a-j\}} \quad . \tag{5}$$

These equations can be worked to obtain the parameters scalings presented in equation (1) in the main text.

### Supplementary Text S2. Expected divergence of PoMos

To assess the impact of selection in branch lengths estimation, we computed the expected divergence per generation for the PoMo models. Unlike the standard models of evolution, which measure divergence in substitutions, the PoMo models measure divergence in terms of frequency shifts. The rate of the process, or divergence $D$, can be formally described as

$$D = -\sum_{i=j} \psi_i q_{ij}$$
$$= \frac{2N \sum_{a_i a_j \in A^C} \pi_{a_i} \rho_{a_i a_j} \pi_{a_j} \sum_{n=1}^{N} \phi_{a_i}^{n-1} \phi_{a_j}^{N-n}}{\sum_{a_i \in A} \pi_{a_i} \phi_{a_i}^{N-1} + \sum_{a_i a_j \in A^C} \pi_{a_i} \pi_{a_j} \rho_{a_i a_j} \sum_{n=1}^{N-1} \phi_{a_i}^{n-1} \phi_{a_j}^{N-n-1} [n\phi_{a_i} + (N-n)\phi_{a_j}] \frac{N}{n(N-n)}}, \quad (6)$$

where $A$ is an alphabet of $K$ alleles and $A^C$ all the possible pair-wise comparisons of $K$ alleles (representing all the edges in the PoMo state space). We adapted equation 6 for the biallelic case; the following code permits to calculate the expected divergence for diverse mutation-selection scenarios. For the biallelic case, the mutational bias can be easily determined by the ratio of the base compositions of the two alleles because $\mu_{Aa}/\mu_{aA} = \pi a/\pi_A$.

```
# function rate of the process
# calculates the expected divergence per generation for the biallelic case
# sigmaA and sigmaa are the selection coefficients
# pia, piA and rhoAa are the parameters of the mutation rates: muAa=pia*rhoAa and muaA=piA*rhoAa
# N is the effective population size

rate_process <- function(N,pia,piA,rhoAa,sigmaa,sigmaA){
  n <- 1:N
  rate <- sum(2*pia*piA*rhoAa*((1+sigmaa)^(n-1))*((1+sigmaA)^(N-n)))
  n <- 1:(N-1)
  norm <- pia*((1+sigmaa)^(N-1))+ piA*((1+sigmaA)^(N-1)) +
          sum( pia*piA*rhoAa*((1+sigmaa)^(n-1))*((1+sigmaA)^(N-n-1))*
                         (n*(1+sigmaa)+(N-1)*(1+sigmaA))*N/(n*(N-n))  )
  return(rate/norm)
}
```

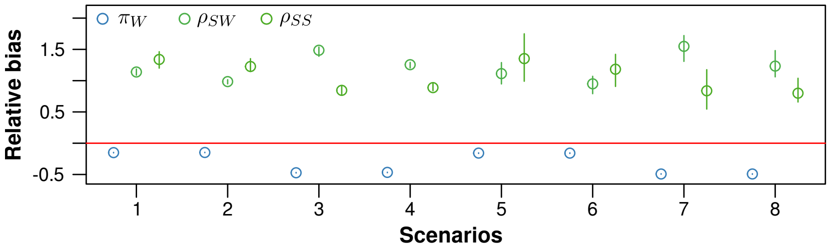


Figure 1: **Estimating mutation rates and selection coefficients with a virtual dynamic.** The simulated data from a four-allelic Moran dynamic with mutation bias and selection was feed to the virtual PoMoTwo and the population parameters estimated using the Bayesian phylogenetic tool RevBayes (Höhna et al., 2016). As the population parameters were estimated in a population of two virtual individuals, we recalibrated them back to the effective population using the scaling laws described in the main text [equations (1) and (2)]: i.e., $\pi_i = \pi_j^*$ and $\rho_{a_i a_j} N H_{N-1} = 2\rho^*_{a_i a_j}$. The estimated parameters are four: $\pi_W$, $\rho_{SW}$ and $\rho_{SS}$ are terms of the mutation rates, that were assumed reversible (i.e., $\mu_{SW} = \rho_{SW}\pi_W$); $\phi_S$ is the fitness coefficient of the strong alleles. The average bias of the estimates was obtained based on alignments of 100000 sites and 25 replicates per scenario.

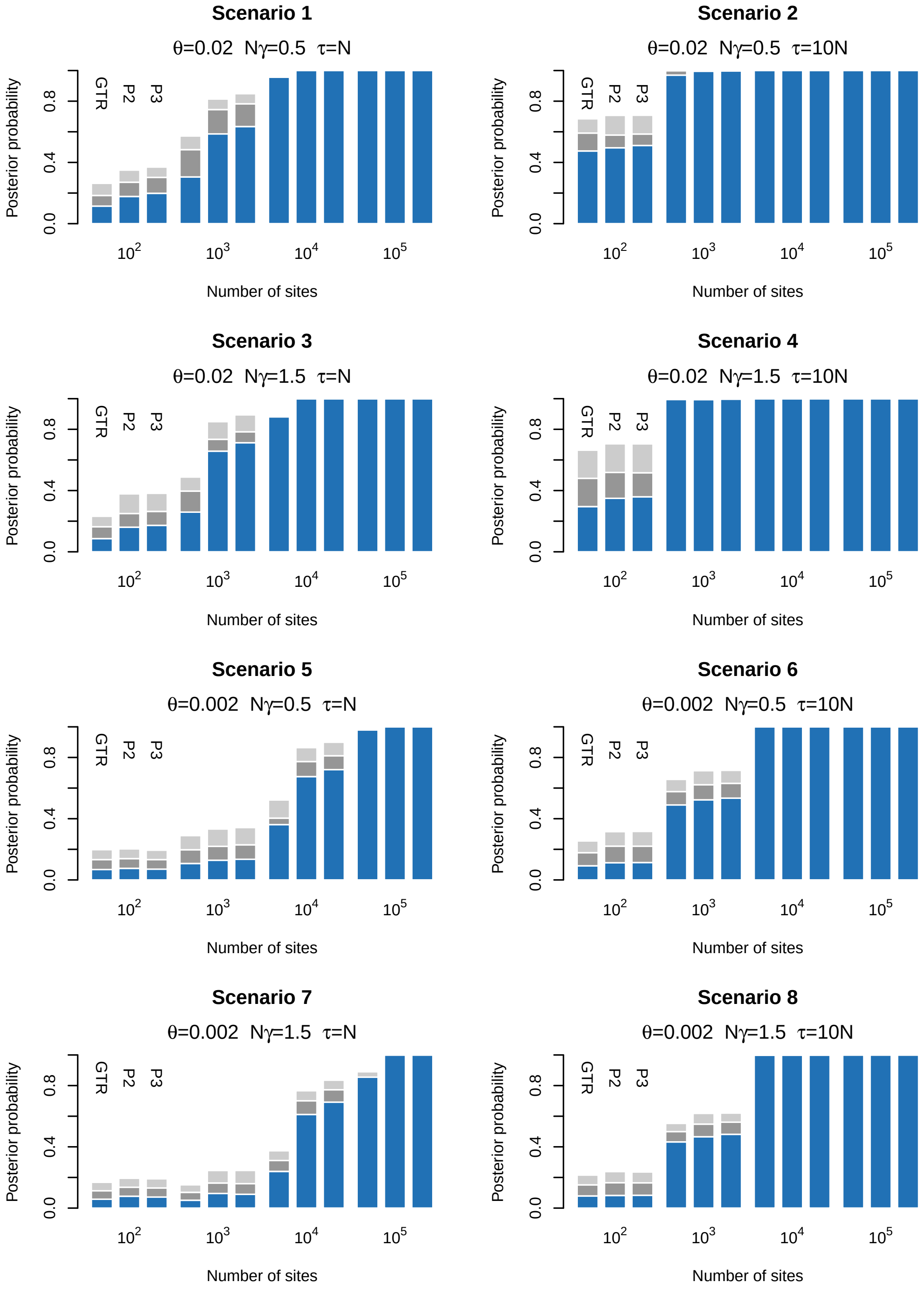


Figure 2: **Estimating the species tree topology with selection.** Data sets of different sizes (x-axis) and simulated under different scenarios 1-8 were used to perform species tree inferences with three methods: GTR, PoMoTwo (P2 in the figure), and PoMoThree (P3 in the figure). We investigated the accuracy of the topology by assessing the posterior probability of the true topology (`(A,(B,(C,D)))`; blue sub-bars) under these three methods. The remaining probability correspond to wrongly estimated topologies. The posterior probabilities of wrong topologies that are still able to correctly identify the first taxum to diverge are represented in shades of gray: `(A,(C,(B,D)))` in dark gray and `(A,(D,(C,B)))` in light gray. PoMoTwo and Three trees inferred based on a sample of 10 individuals per taxa (i.e., 40 sequences in total).

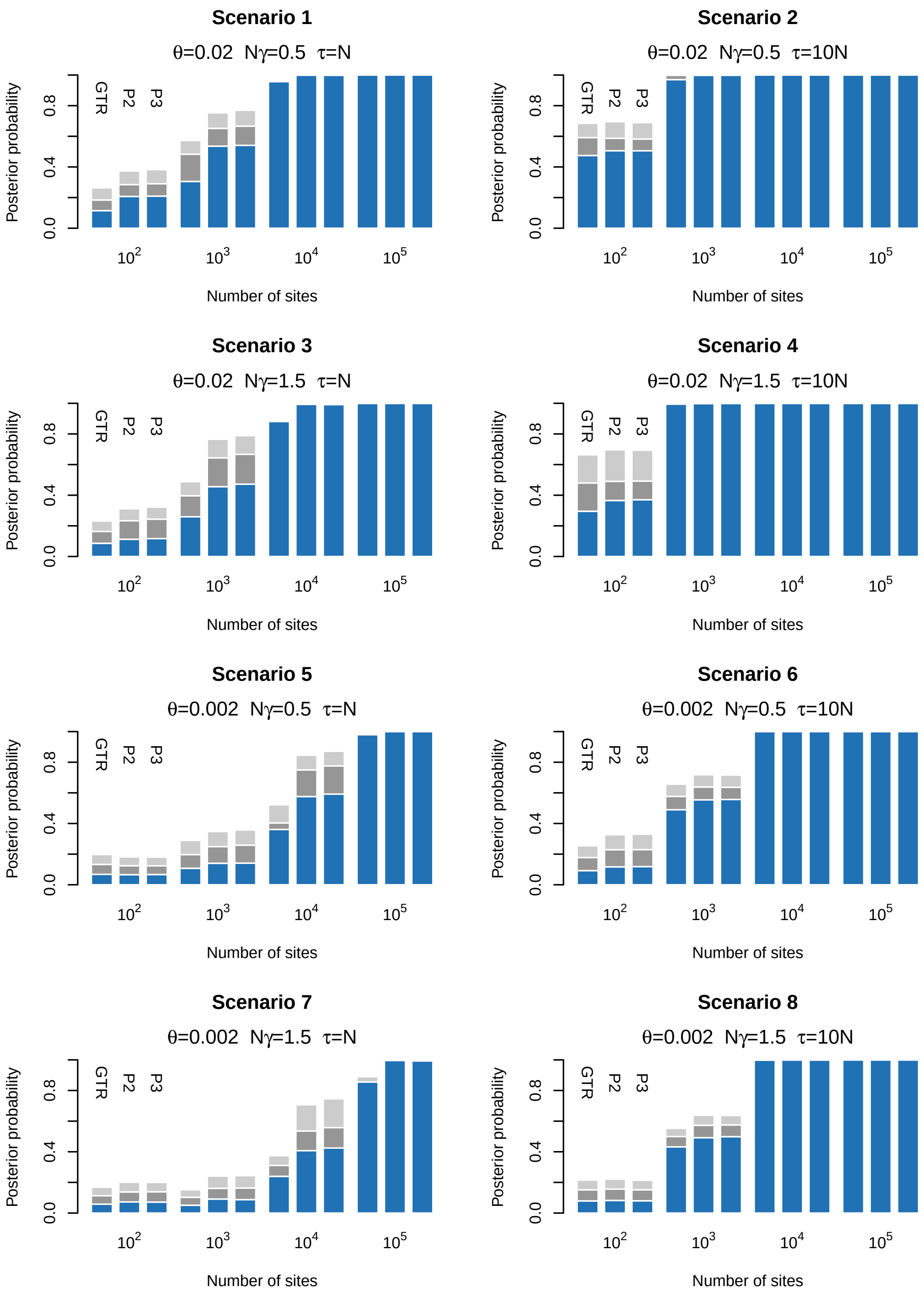


Figure 3: **Estimating the species tree topology with selection.** Data sets of different sizes (x-axis) and simulated under different scenarios 1-8 were used to perform species tree inferences with three methods: GTR, PoMoTwo (P2 in the figure), and PoMoThree (P3 in the figure). We investigated the accuracy of the topology by assessing the posterior probability of the true topology (`(A,(B,(C,D)))`; blue sub-bars) under these three methods. The remaining probability correspond to wrongly estimated topologies. The posterior probabilities of wrong topologies that are still able to correctly identify the first taxum to diverge are represented in shades of gray: `(A,(C,(B,D)))` in dark gray and `(A,(D,(C,B)))` in light gray. PoMoTwo and Three trees inferred based on a sample of 2 individuals per taxa (i.e., 8 sequences in total).

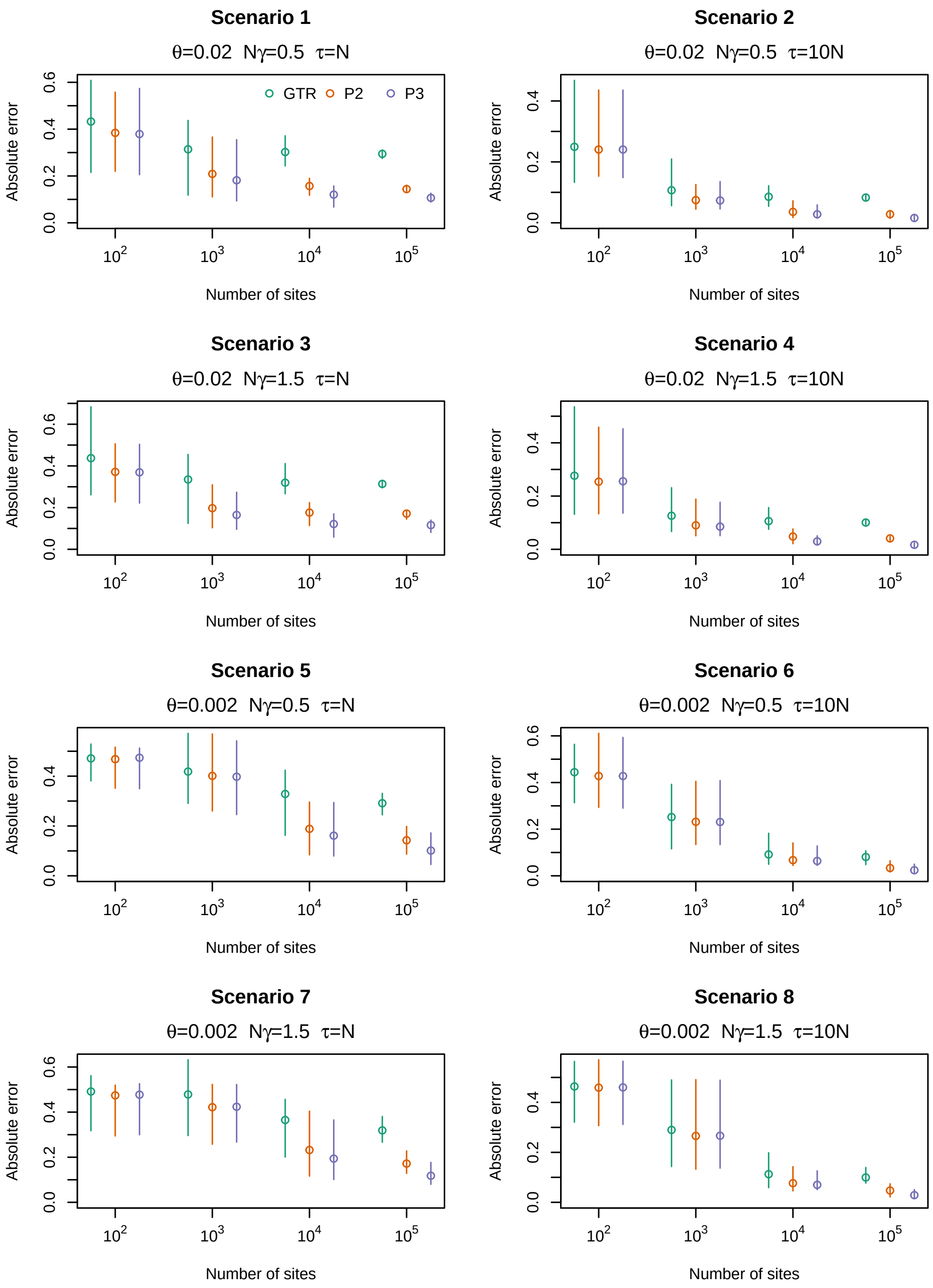


Figure 4: **Estimating the species tree branch lenghts with selection.** Data sets of different sizes (x-axis) and simulated under different scenarios 1-8 were used to perform species tree inferences with three methods: GTR, PoMoTwo (P2 in the figure), and PoMoThree (P3 in the figure). We investigated the accuracy of the branch lengths by calculating the absolute error of all six pairwise comparisons of genetic distances among the four taxa. As the three methods employ different evolutionary units, the branch lengths were normalized so that the three models could be compared. PoMoTwo and Three trees inferred based on a sample of 10 individuals per taxa (40 sequences in total).

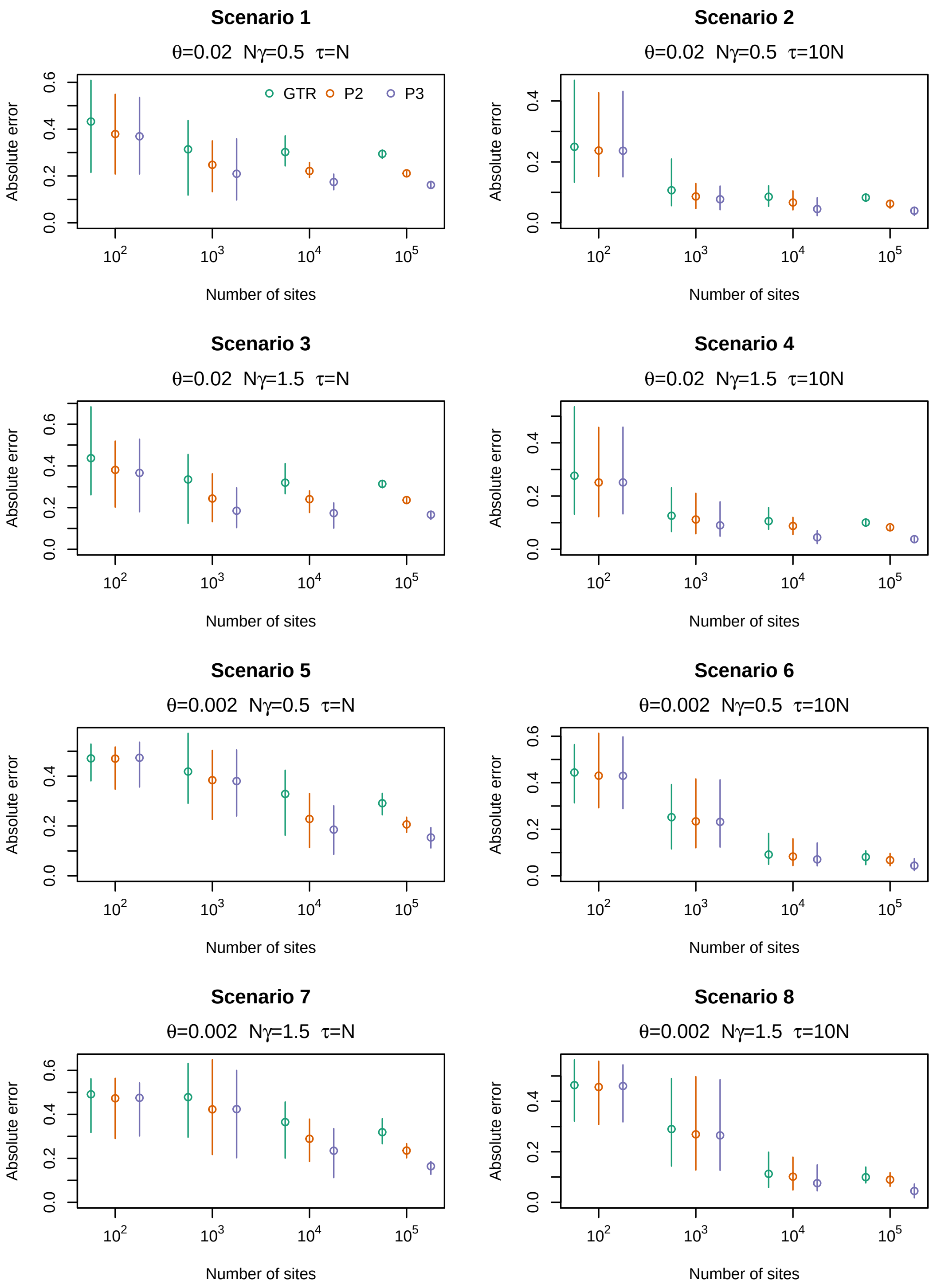


Figure 5: **Estimating the species tree branch lenghts with selection.** Data sets of different sizes (x-axis) and simulated under different scenarios 1-8 were used to perform species tree inferences with three methods: GTR, PoMoTwo (P2 in the figure), and PoMoThree (P3 in the figure). We investigated the accuracy of the branch lengths by calculating the absolute error of all six pairwise comparisons of genetic distances among the four taxa. As the three methods employ different evolutionary units, the branch lengths were normalized so that the three models could be compared. PoMoTwo and Three trees inferred based on a sample of 2 individuals per taxa (8 sequences in total).

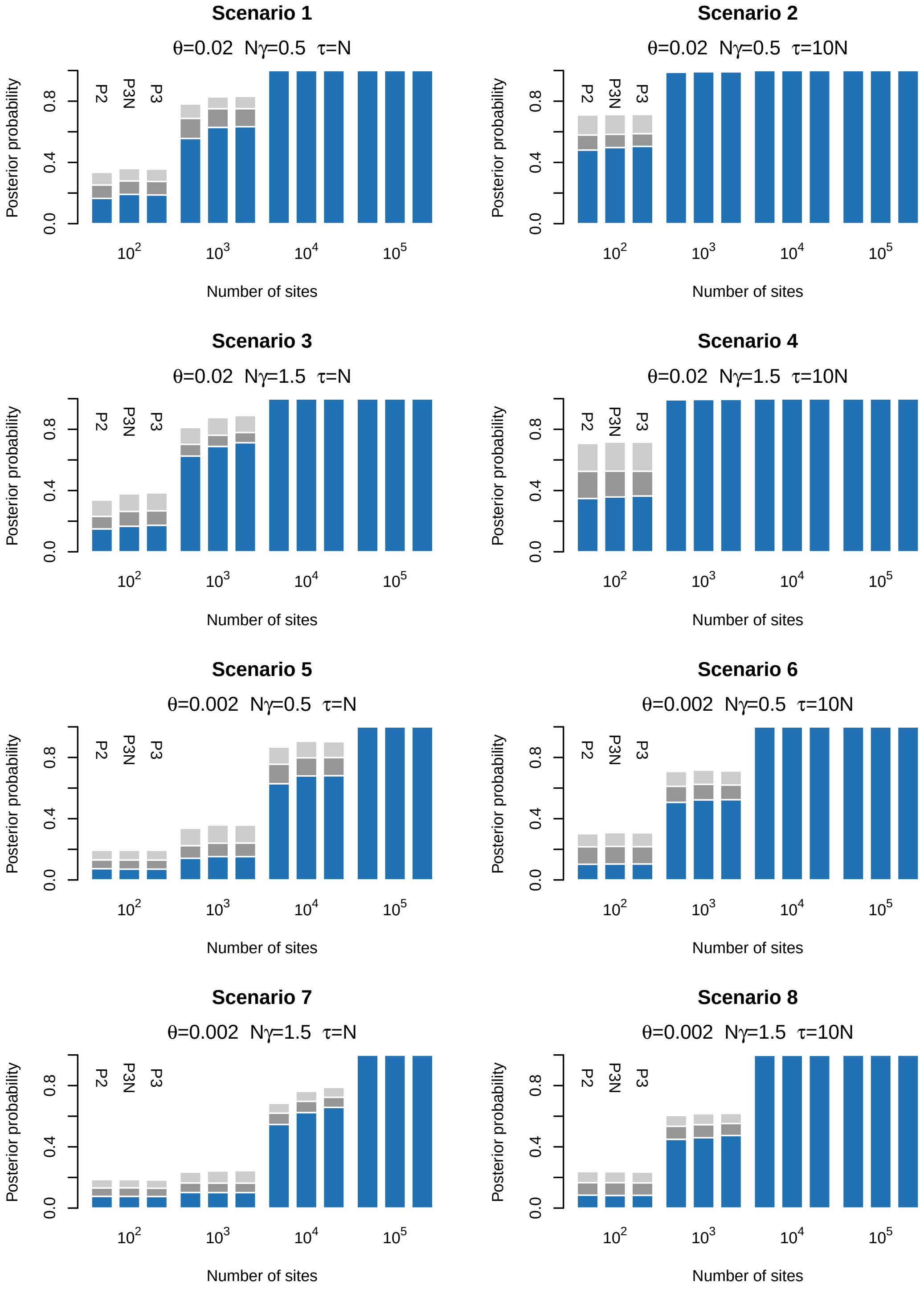


Figure 6: **Estimating the species tree topology with selection.** Data sets of different sizes (x-axis) and simulated under different scenarios 1-8 were used to perform species tree inferences with three methods: PoMoTwo (P2 in the figure), and PoMoThree with fitness coefficients fixed to 1.0 (P3N in the figure) and PoMoThree (P3 in the figure). We investigated the accuracy of the topology by assessing the posterior probability of the true topology (`(A,(B,(C,D)))`; blue sub-bars) under these three methods. The remaining probability correspond to wrongly estimated topologies. The posterior probabilities of wrong topologies that are still able to correctly identify the first taxum to diverge are represented in shades of gray: `(A,(C,(B,D)))` in dark gray and `(A,(D,(C,B)))` in light gray. PoMoTwo and Three trees inferred based on a sample of 10 individuals per taxa (i.e., 40 sequences in total).

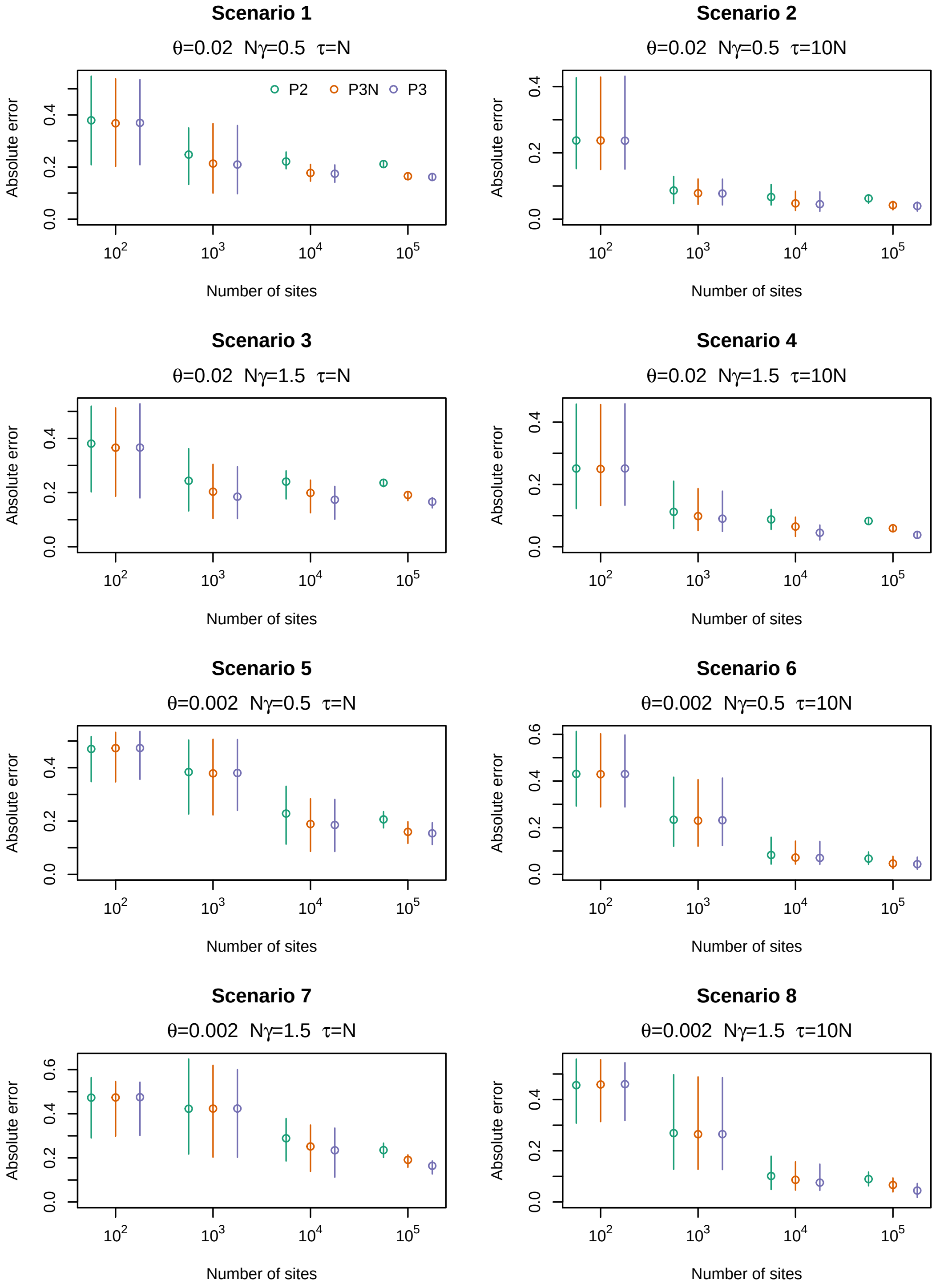


Figure 7: **Estimating the species tree branch lenghts with selection.** Data sets of different sizes (x-axis) and simulated under different scenarios 1-8 were used to perform species tree inferences with three methods: PoMoTwo (P2 in the figure), and PoMoThree with fitness coefficients fixed to 1.0 (P3N in the figure) and PoMoThree (P3 in the figure). We investigated the accuracy of the branch lengths by calculating the absolute error of all six pairwise comparisons of genetic distances among the four taxa. As the three methods employ different evolutionary units, the branch lengths were normalized so that the three models could be compared. PoMoTwo and Three trees inferred based on a sample of 10 individuals per taxa (40 sequences in total).

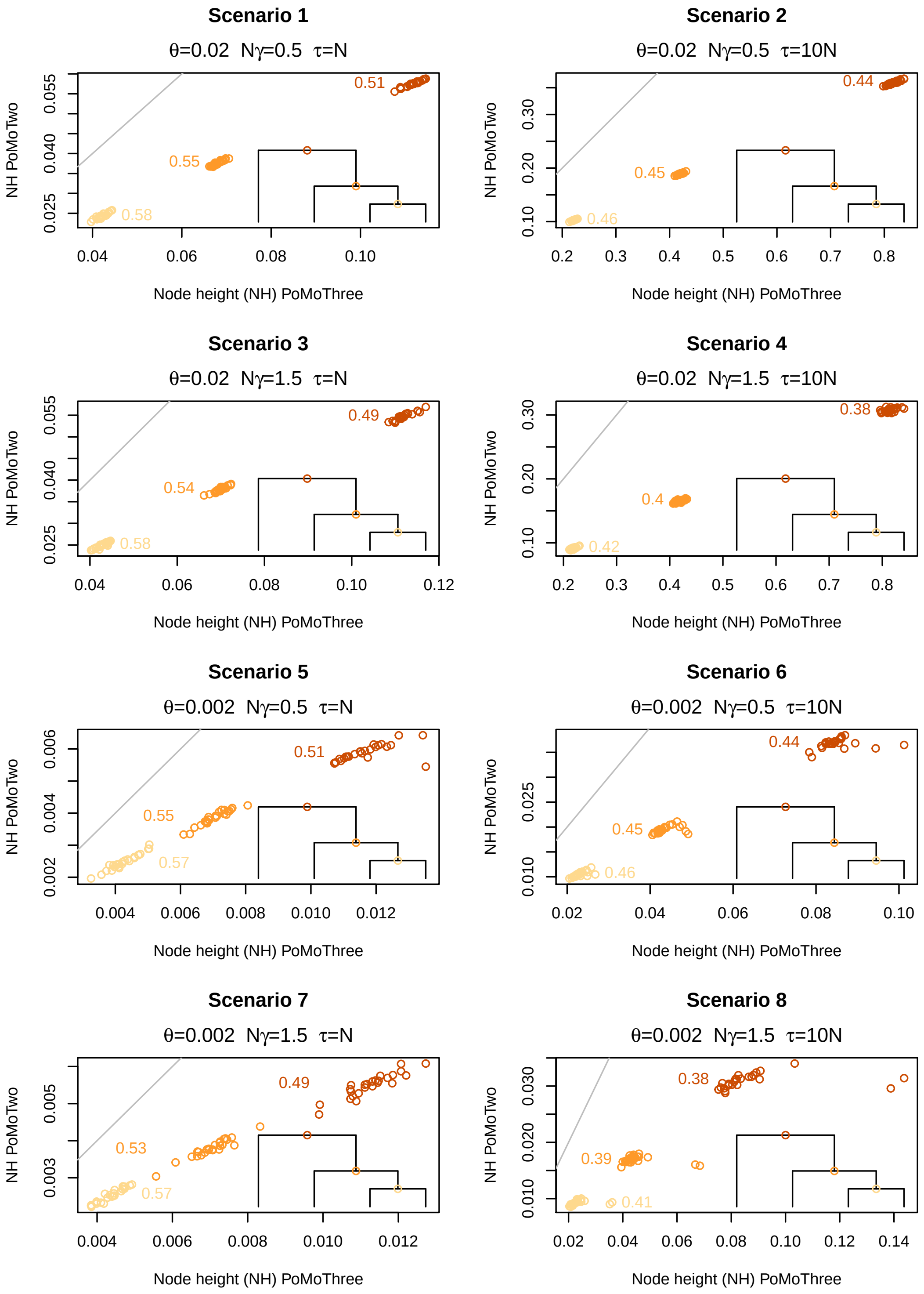


Figure 8: **Node height estimation under the virtual PoMos.** The phylogenetic tree depicts the three nodes' placement in the true phylogeny: the node heights double as we approach the root. Values accompanying each cloud denote the average ratio of the node heights in PoMoTwo to those in PoMoThree. Each cloud represents 25 replicated alignments. These distances were estimated using only the 100 000 sites alignments, where both methods return the true topology with maximum accuracy. Node heights inferred based on a sample of 10 individuals per taxa (40 sequences in total)

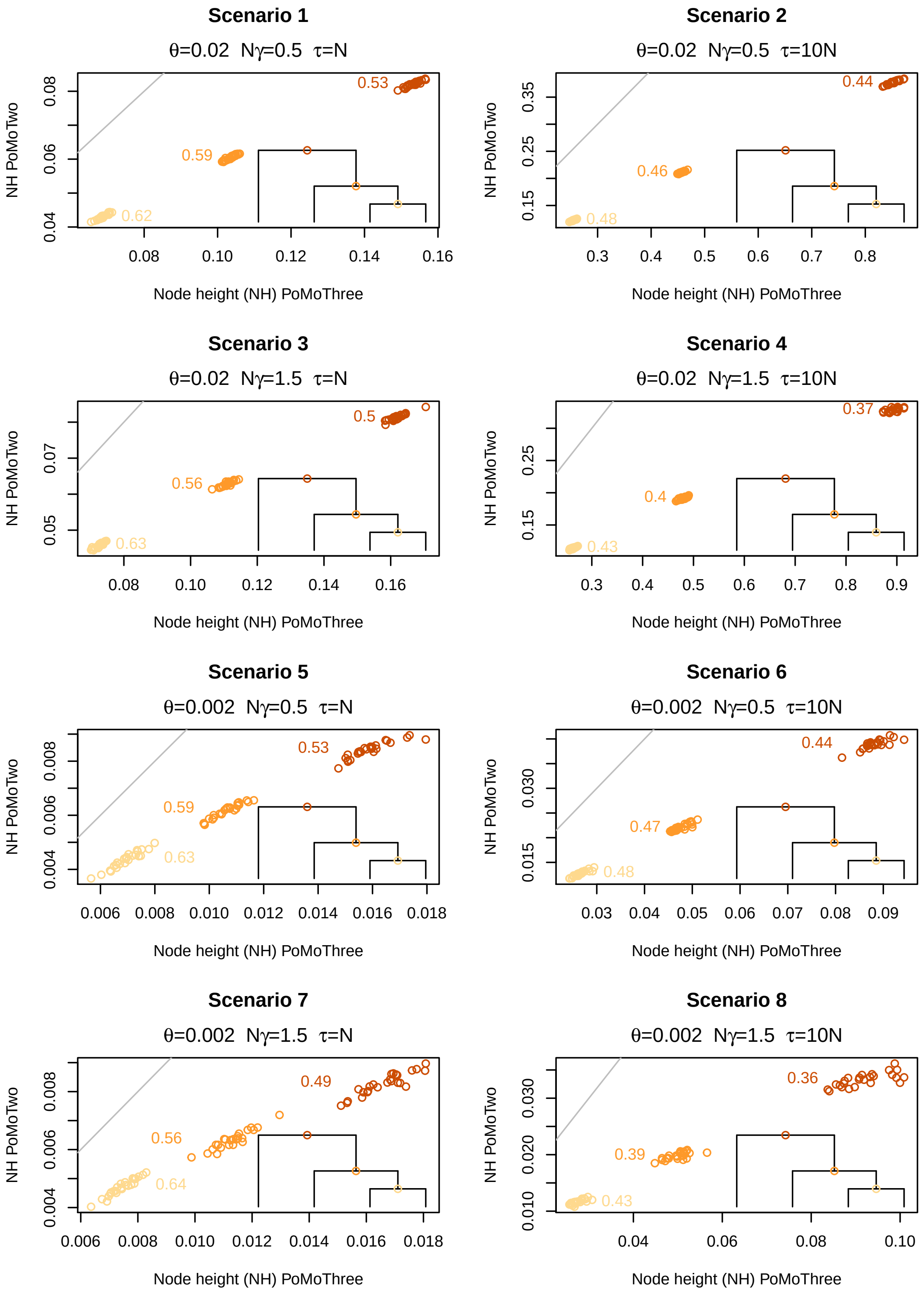


Figure 9: **Node height estimation under the virtual PoMos.** The phylogenetic tree depicts the three nodes' placement in the true phylogeny: the node heights double as we approach the root. Values accompanying each cloud denote the average ratio of the node heights in PoMoTwo to those in PoMoThree. Each cloud represents 25 replicated alignments. These distances were estimated using only the 100 000 sites alignments, where both methods return the true topology with maximum accuracy. Node heights inferred based on a sample of 2 individuals per taxa (8 sequences in total)

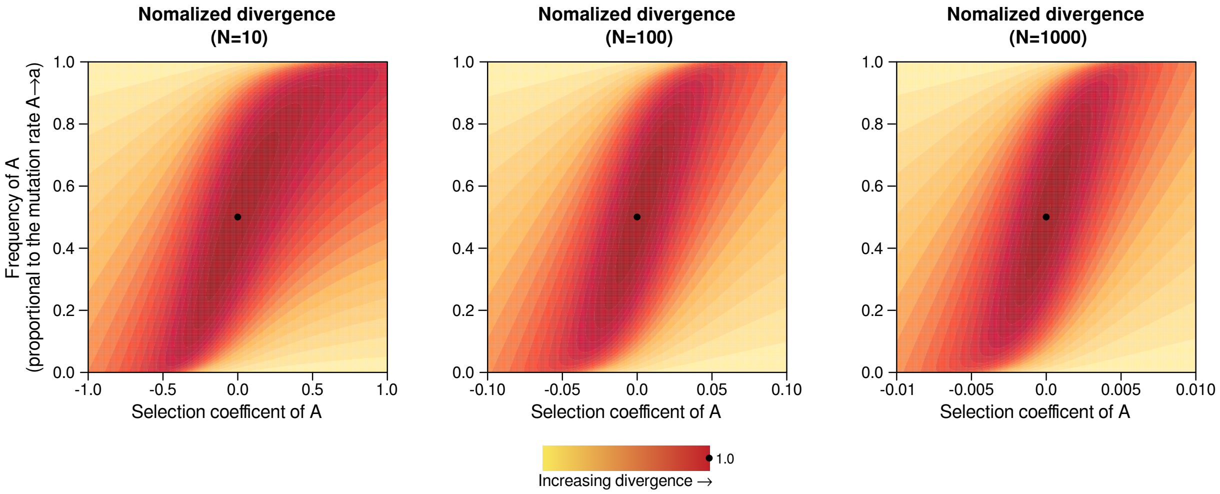


Figure 10: **Impact of the effective population size on the expected divergence.** The expected divergence was calculated for different regimes of selection and mutation and normalized by the middle point (no selection nor mutational bias). For the biallelic case, the base composition $\pi$ is proportional to the mutational bias: $\frac{\mu_{Aa}}{\mu_{aA}} = \frac{\pi_a}{\pi_A}$, where $\mu$ is the mutation rate. If the base composition is equal ($y = 0.5$), there is no mutational bias.

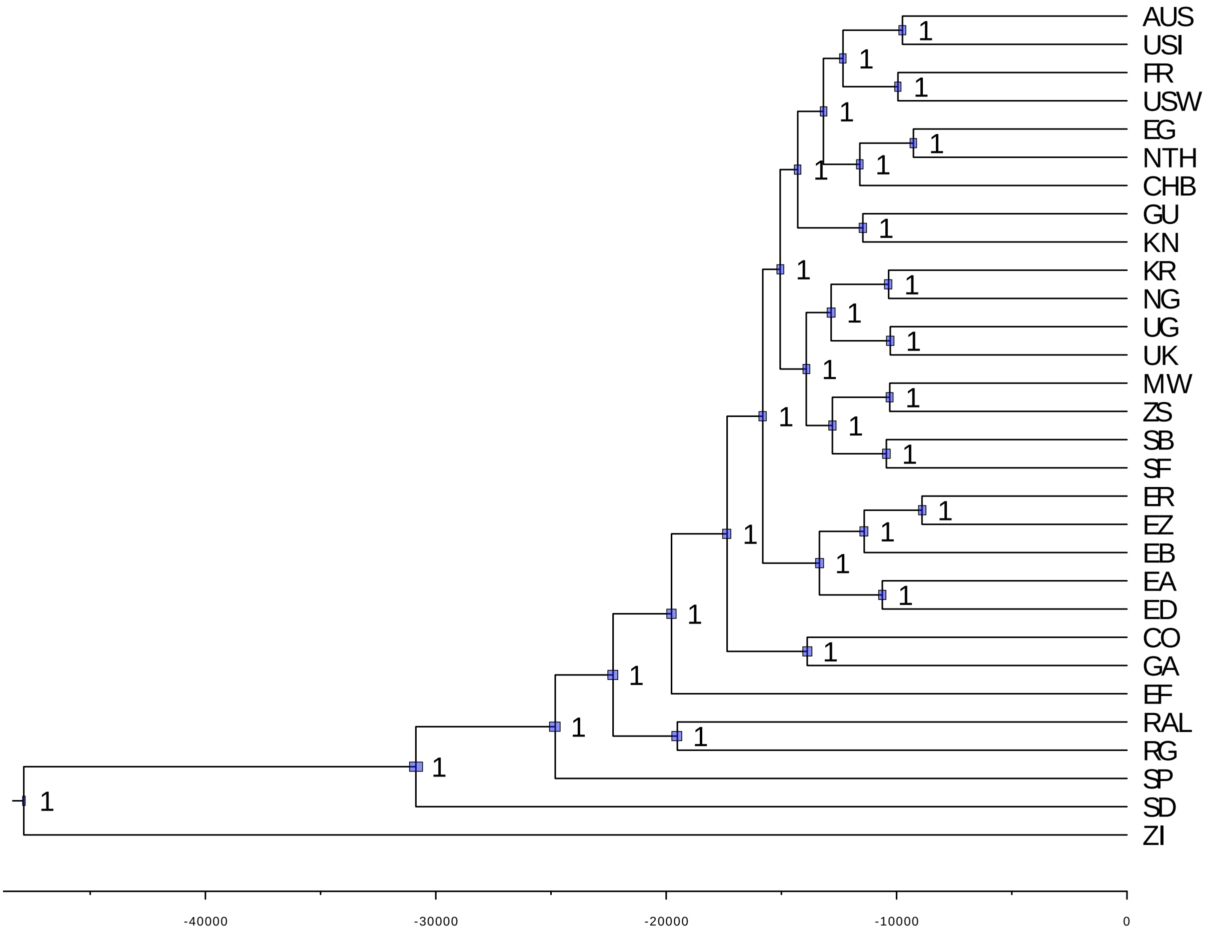


Figure 11: **Molecular dating of the fruit flies' evolutionary history with the GTR model.** Phylogenetic inferences were performed in RevBayes together with the GTR model of evolution and by employing a Markov chain Monte Carlo scheme with two chains and 150 000 iterates. The depicted phylogeny corresponds to the maximum a posteriori tree. Population location: AUS (Australia, Sorell TAS), CHB (China, Beijing), CO (Cameroon, Oku), EA (Ethiopia, Gambella), EB (Ethiopia, Bonga), ED (Ethiopia, Dodola), EF (Ethiopia, Fiche), EG (Egypt, Cairo), ER (Ethiopia, Debre Birhan), EZ (Ethiopia, Ziway), FR (France, Lyon), GA (Gabon, Franceville), GU (Guinea, Donde), KN (Kenya, Nyahururu), KR (Kenya, Marigat), MW (Malawi), NG (Nigeria, Maiduguri), NTH (Netherlands, Houten), RAL (United States, Raleigh NC), RG (Rwanda, Gikongoro), SB (South Africa, Barkly East), SD (South Africa, Dullstroom), SF (South Africa, Fouriesburg), SP (South Africa, Phalaborwa), UG (Uganda, Namulonge), UK (Uganda, Kisoro), USI (United States, Ithaca NY), USW (United States, Winters CA), ZI (Zambia, Siavonga), ZS (Zimbabwe, Sengwa).

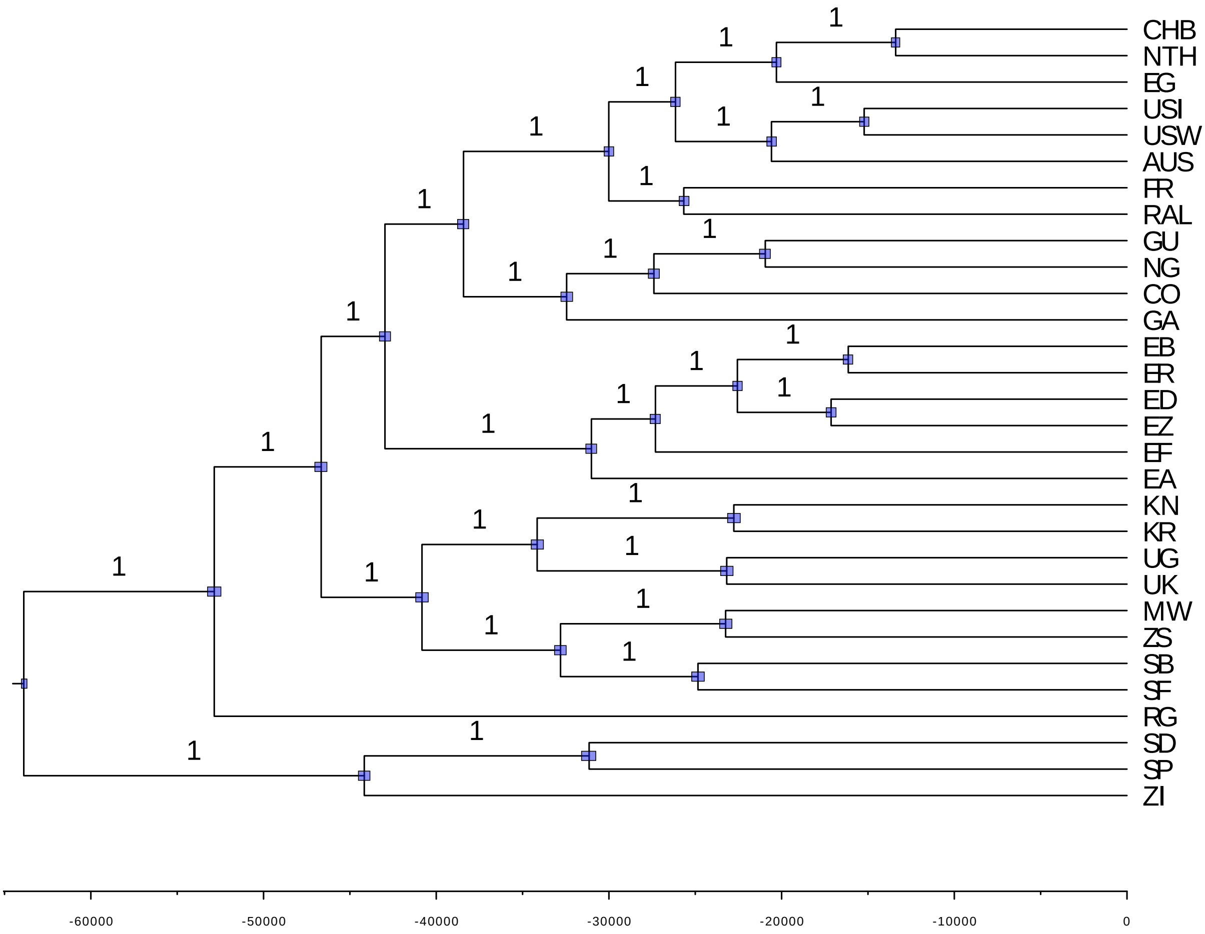


Figure 12: **Molecular dating of the fruit flies' evolutionary history with the PoMoTwo model.** Phylogenetic inferences were performed in RevBayes together with the PoMoTwo model of evolution and by employing a Markov chain Monte Carlo scheme with two chains and 150 000 iterates. The depicted phylogeny corresponds to the maximum a posteriori tree. Population location: AUS (Australia, Sorell TAS), CHB (China, Beijing), CO (Cameroon, Oku), EA (Ethiopia, Gambella), EB (Ethiopia, Bonga), ED (Ethiopia, Dodola), EF (Ethiopia, Fiche), EG (Egypt, Cairo), ER (Ethiopia, Debre Birhan), EZ (Ethiopia, Ziway), FR (France, Lyon), GA (Gabon, Franceville), GU (Guinea, Donde), KN (Kenya, Nyahururu), KR (Kenya, Marigat), MW (Malawi), NG (Nigeria, Maiduguri), NTH (Netherlands, Houten), RAL (United States, Raleigh NC), RG (Rwanda, Gikongoro), SB (South Africa, Barkly East), SD (South Africa, Dullstroom), SF (South Africa, Fouriesburg), SP (South Africa, Phalaborwa), UG (Uganda, Namulonge), UK (Uganda, Kisoro), USI (United States, Ithaca NY), USW (United States, Winters CA), ZI (Zambia, Siavonga), ZS (Zimbabwe, Sengwa)

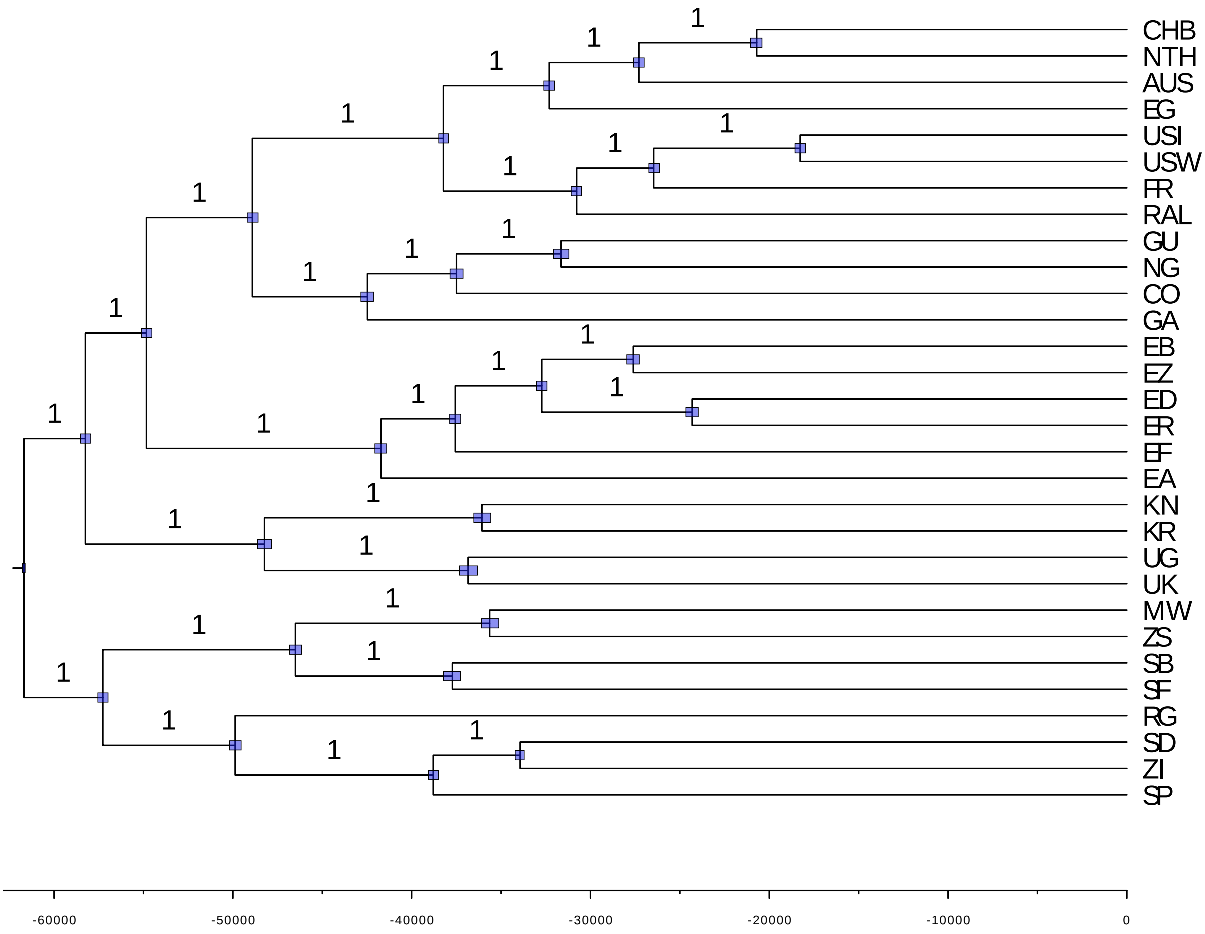


Figure 13: **Molecular dating of the fruit flies' evolutionary history with the PoMoThree model.** Phylogenetic inferences were performed in RevBayes together with the PoMoThree model of evolution and by employing a Markov chain Monte Carlo scheme with two chains and 150 000 iterates. The depicted phylogeny corresponds to the maximum a posteriori tree. Population location: AUS (Australia, Sorell TAS), CHB (China, Beijing), CO (Cameroon, Oku), EA (Ethiopia, Gambella), EB (Ethiopia, Bonga), ED (Ethiopia, Dodola), EF (Ethiopia, Fiche), EG (Egypt, Cairo), ER (Ethiopia, Debre Birhan), EZ (Ethiopia, Ziway), FR (France, Lyon), GA (Gabon, Franceville), GU (Guinea, Donde), KN (Kenya, Nyahururu), KR (Kenya, Marigat), MW (Malawi), NG (Nigeria, Maiduguri), NTH (Netherlands, Houten), RAL (United States, Raleigh NC), RG (Rwanda, Gikongoro), SB (South Africa, Barkly East), SD (South Africa, Dullstroom), SF (South Africa, Fouriesburg), SP (South Africa, Phalaborwa), UG (Uganda, Namulonge), UK (Uganda, Kisoro), USI (United States, Ithaca NY), USW (United States, Winters CA), ZI (Zambia, Siavonga), ZS (Zimbabwe, Sengwa)

| No. of species | No. of sites | GTR | PoMoTwo | PoMoThree |
|---|---|---|---|---|
| 10 | 100 | 2050 | 1851 | 1608 |
| 20 | 100 | 3342 | 3218 | 2219 |
| 50 | 100 | 4179 | 1113 | 2570 |
| 100 | 100 | 3684 | 2316 | 2974 |
| 10 | 1000 | 2234 | 2691 | 2557 |
| 20 | 1000 | 3656 | 3046 | 983 |
| 50 | 1000 | 3790 | 3495 | 1975 |
| 100 | 1000 | 2433 | 3833 | 3066 |
| 10 | 10000 | 4030 | 3086 | 506 |
| 20 | 10000 | 2224 | 3832 | 2340 |
| 50 | 10000 | 3090 | 2529 | 3508 |
| 100 | 10000 | 1661 | 4197 | 2871 |
| 10 | 100000 | 3884 | 2645 | 2995 |
| 20 | 100000 | 3474 | 2558 | 1522 |
| 50 | 100000 | 4189 | 3935 | 3063 |
| 100 | 100000 | 4708 | 4465 | 2798 |
| 10 | 1000000 | 1944 | 2399 | 1494 |
| 20 | 1000000 | 2281 | 3348 | 1470 |
| 50 | 1000000 | 4286 | 3775 | * |
| 100 | 1000000 | * | * | * |

Table 1: **Average Effective Sample Size of the sample posterior parameter under the GTR and virtual PoMoTwo and Three models.** The Bayesian inferences consisted of a standard MCMC chain with 50 000 generations and on simulated alignments with 5 to 100 species and 100 to 1 million sites with levels of diversity similar to those of great apes. These analyses ran on an iMac desktop with an Intel Core i5 processor of 3.4 GHz of speed and 32GB of memory, without any kind of parallelization. * The analyses did not started because the used memory reached the computers' maximum.

| Population | Country | Locality | Samples |
|---|---|---|---|
| AUS | Australia | Sorell TAS | 18 |
| CHB | China | Beijing | 15 |
| CO | Cameroon | Oku | 10 |
| EA | Ethiopia | Gambella | 24 |
| EB | Ethiopia | Bonga | 5 |
| ED | Ethiopia | Dodola | 5 |
| EF | Ethiopia | Fiche | 69 |
| EG | Egypt | Cairo | 32 |
| ER | Ethiopia | Debre Birhan | 5 |
| EZ | Ethiopia | Ziway | 4 |
| FR | France | Lyon | 96 |
| GA | Gabon | Franceville | 10 |
| GU | Guinea | Donde | 5 |
| KN | Kenya | Nyahururu | 5 |
| KR | Kenya | Marigat | 4 |
| MW | Malawi | - | 9 |
| NG | Nigeria | Maiduguri | 6 |
| NTH | Netherlands | Houten | 19 |
| RAL | United States | Raleigh NC | 205 |
| RG | Rwanda | Gikongoro | 27 |
| SB | South Africa | Barkly East | 5 |
| SD | South Africa | Dullstroom | 81 |
| SF | South Africa | Fouriesburg | 5 |
| SP | South Africa | Phalaborwa | 37 |
| UG | Uganda | Namulonge | 4 |
| UK | Uganda | Kisoro | 5 |
| USI | United States | Ithaca NY | 19 |
| USW | United States | Winters CA | 35 |
| ZI | Zambia | Siavonga | 197 |
| ZS | Zimbabwe | Sengwa | 5 |
| **Total** | | | 966 |

Table 2: **Fruit fly genomic data sets used for the molecular dating analyses.** The data was taken from FlyPop (Hervas et al., 2017).
